# Transcriptomic signatures of sex-specific nicotine sensitization and imprinting of self-administration in rats inform GWAS findings on human addiction phenotypes

**DOI:** 10.1101/2020.10.05.327155

**Authors:** Alena Kozlova, Robert R Butler, Siwei Zhang, Thomas Ujas, Hanwen Zhang, Stephan Steidl, Alan R. Sanders, Zhiping P. Pang, Paul Vezina, Jubao Duan

## Abstract

Rodents are frequently used to model drug addiction, yet their genetic relevance to human addictive behaviors especially the mounting genome-wide association study (GWAS) findings is poorly understood. Considering a possible gateway drug role of nicotine (NIC), we modeled NIC addiction, specifically NIC sensitization (SST) and self-administration (SA), in F1 progeny of inbred Envigo rats (F344/BN) and conducted integrative genomics analyses. We unexpectedly observed male-specific NIC SST and a parental effect of SA only present in paternal F344 crosses. Transcriptional profiling in the ventral tegmental area (VTA) and nucleus accumbens (NAc) core and shell further revealed sex and brain region-specific transcriptomic signatures of SST and SA. We found that genes associated with SST and SA were enriched for those related to synaptic processes, myelin sheath, and tobacco use disorder or chemdependency. Interestingly, SST-associated genes were often downregulated in male VTA but upregulated in female VTA, and strongly enriched for smoking GWAS risk variants, possibly explaining the male-specific SST. For SA, we found widespread region-specific allelic imbalance of expression (AIE), of which genes showing AIE bias towards paternal F344 alleles in NAc core were strongly enriched for SA-associated genes and for GWAS risk variants of smoking initiation, likely contributing to the parental effect of SA. The transcriptional signatures of sex-specific nicotine SST and SA suggest a mechanistic link between genes underlying these processes and human nicotine addiction, providing a resource for understanding the biology underlying the GWAS findings on human smoking and other addictive phenotypes.

## Introduction

Cigarette smoking is the most common contributor to premature death worldwide. It is also a major risk factor for many diseases, including cancer ^1^ and cardiovascular diseases ^2^. Among over 4,800 chemical compounds in tobacco, nicotine (NIC) determines the addictive nature of smoking and is the principal psychoactive ingredient of tobacco products ^3^. NIC is also a possible gateway “drug” for other substances of abuse^4, 5^. NIC addiction in humans is a polygenic disorder. Genome-wide association studies (GWAS) of cigarette smoking have identified a plethora of genetic loci with common single nucleotide polymorphisms (SNPs) associated with different smoking phenotypes ^6-10^. In the most recent GWAS meta-analysis with ∼1.2 million individuals, genetic variants at 406 loci were found associated with smoking initiation (SIn), cessation (SCe), cigarettes per day (CPD), and age of initiation (AOI) ^11^. Consistent with the known biology of NIC addiction, the dopamine D2 receptor (DRD2) gene and almost all central nervous system (CNS)-expressed nicotinic acetylcholine receptor (nAChR) genes are significantly associated with one or more smoking phenotypes ^11^. However, the molecular mechanisms underlying most smoking GWAS risk loci remain elusive. Furthermore, because each smoking locus often spans many equivalently associated genetic variants and multiple genes, the exact risk genes in many of these GWAS risk loci remain undetermined.

Much of our knowledge of NIC function stems from the study of NIC addiction models in rodents. NIC, like many other abused substances such as amphetamine and cocaine, activates brain dopaminergic neurotransmission and increases locomotor activity in laboratory animals ^12^. A key process of developing NIC-related addictive behaviors in rodents is NIC sensitization (SST). SST reflects long-lasting non-associative neuroadaptations whereby the ability of drugs (such as NIC) to activate dopamine (DA) neurotransmission and elicit appetitive behaviors is enhanced ^4, 5^. When NIC is repeatedly administered, its effects are enhanced so that re-exposure to the drug, weeks to months later in rodents, produces greater behavioral activation (i.e., locomotor activity) than was initially observed ^4, 5^. NIC SST may have relevance to the initiation, maintenance, and escalation of NIC use that is characteristic of the transition from casual experimentation with NIC or other addictive drugs to craving and abuse in humans ^4^. Although it is commonly known that nAChRs expressed in the ventral tegmental area (VTA) mediate the ability of NIC to increase locomotion and DA release in the nucleus accumbens (NAc) ^13-15^, the SST-associated genome-wide gene expression (i.e., transcriptome) changes and their relevance to genetic findings for human smoking traits remain elusive.

There is also a similar knowledge gap for another commonly studied addiction model, NIC self-administration (SA) in rodents ^3^. Intravenous SA of NIC can be used to determine how motivated animals are to obtain NIC, as indexed by the number of lever presses they are willing to invest in order to earn an infusion ^4, 16^. Prior exposure to NIC leads to enhanced locomotor responding to and SA of the drug in rats ^4, 15, 16^ as well as enhanced cocaine-induced locomotion in C57BL/6J mice ^5^ and amphetamine SA in rats ^17^. However, the molecular link between NIC SST and NIC SA remains unclear, highlighting the need of parallel transcriptomic profiling of NIC SST and SA in relevant brain regions. Furthermore, it remains largely unknown how the gene expression changes associated with SA are relevant to genetic findings of human smoking traits and other addictive behaviors.

Previous gene expression studies of NIC effects in rodents have shown that NIC treatment induces changes in gene expression related to neurological disease pathways, such as G-protein-coupled protein receptor signaling ^18^, glycerolipid metabolism, PI3K and MAPK signaling ^19^, and Hedgehog and Notch signaling pathways ^20^. However, most of these studies assay the acute or direct effects of NIC rather than dissecting the molecular basis of NIC addiction behaviors. Here, we conducted transcriptomic profiling of NIC SST and SA in addiction-relevant rat brain regions: VTA, NAc core and shell. We examined possible sex differences in NIC SST and their possible underlying genetic mechanisms. For the observed parental effect (or imprinting) in NIC SA, we performed allelic imbalance of expression (AIE) analyses to identify genes that may be responsible for this effect. We also demonstrate a potential mechanistic molecular link between NIC SST and SA. The findings have important implications for understanding the genetic mechanisms underlying NIC addiction and help narrow down the risk genes among multiple equally associated genes at each human smoking GWAS risk locus.

## Results

### Modeling NIC SST and SA in F344/BN F1 rats

We modeled NIC SST and SA using F1 progeny of two inbred Envigo strains (Fischer-344 [F344] and Brown Norway [BN]) (Figure 1a, b). Both initial (F1i) and reciprocal crosses (F1r) were carried out to examine possible parent-of-origin effects ^21-23^. The F1i progeny with F344 fathers were labelled as subgroup A, and the F1r progeny with F344 mothers were labelled as subgroup B. The F1 rats are expected to be genetically identical, mitigating the possible confounding effects from individual backgrounds on behavioral and transcriptional analyses. Furthermore, all the F1 rats of the two inbred strains were heterozygous, making them informative for AIE analysis to assess parental effects (Figure 1).

**Figure 1.**
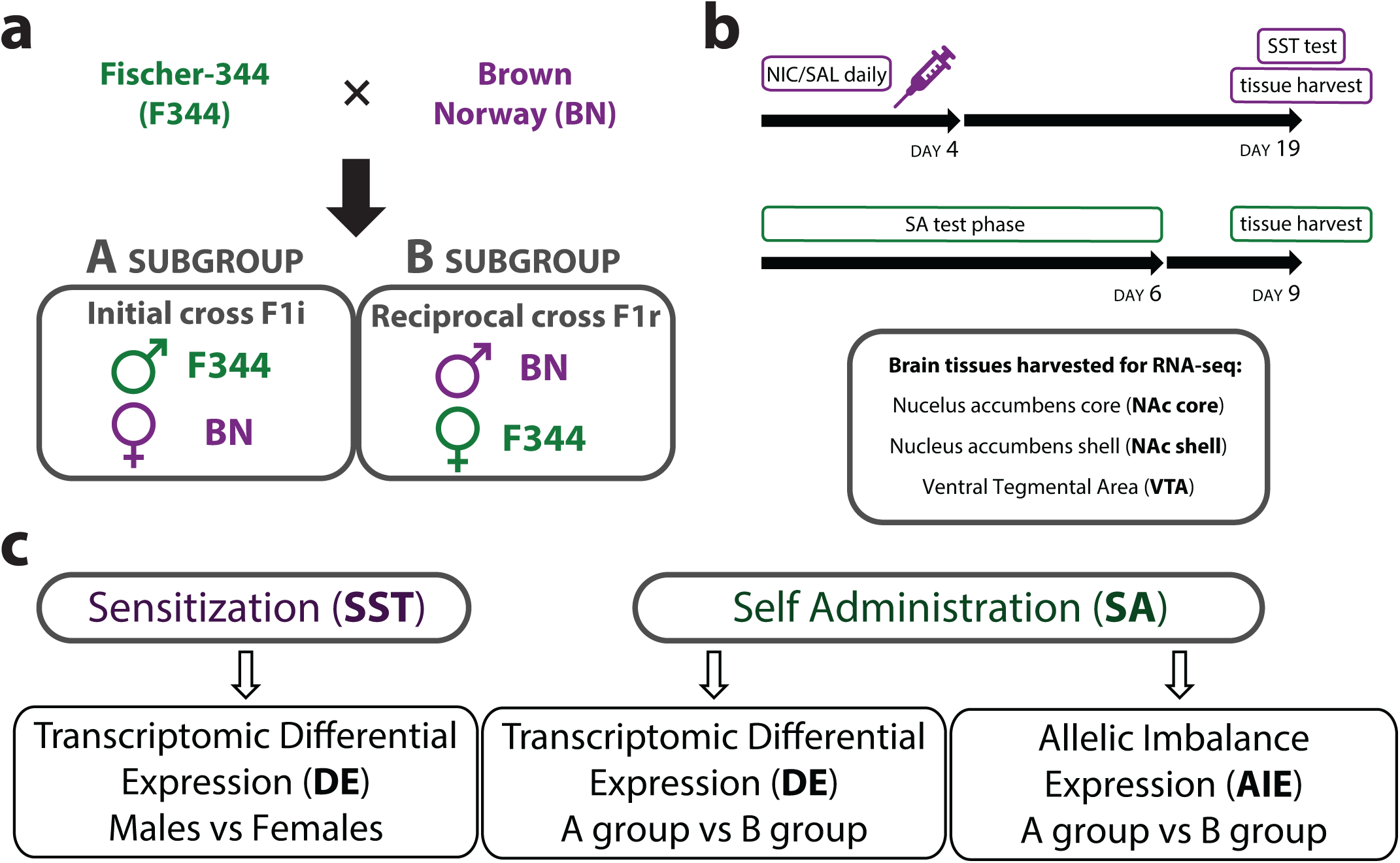
A schematic experimental design for the NIC addiction model. **(a)** Genetically identical and heterozygous F1 progeny of two inbred strains (F344 and BN) were used. To control for parent-of-origin effects, both initial (F1i) and reciprocal (F1r) crosses were carried out to generate F1 rats. “Subgroup A” rats were generated with BN was a mother and “Subgroup B” F1 rats were generated with BN was a father. **(b)** For SST the rats were injected with NIC every day for four days and then tested for SST at day 19. For NIC SA, F1 males were prepared with an IV catheter and given the opportunity to self-administer NIC. For SST transcriptomic analysis, brain tissues from the NAc core, shell and VTA were harvested without testing SST at day 19. For the SA transcriptomic analysis, brain tissues from the same regions were harvested 3 days following the last session. **(c)** Transcriptomic and AIE analyses were performed to identify SST-associated transcriptomic changes between NIC and SAL exposed rats and SA-associated transcriptomic changes in three brain tissues (NAc core, shell and VTA).

NIC SST was tested separately in F1 males (n=5-6/group) and females (n=7/group). Rats in separate groups were administered 0.0 (SAL), 0.1, or 0.4 mg/kg of NIC (base, IP) on each of 4 days. Reciprocal crossed rats (F1i and F1r) were evenly distributed in each group. Locomotor activity was measured on days 1 and 4. Two weeks later, all rats were tested for locomotor SST following a NIC injection (0.4 mg/kg). Additional rats in separate groups were similarly exposed to SAL or NIC (0.4 mg/kg) and sacrificed two weeks later for tissue harvest for RNA-seq analysis (Figure 1b). Relative to SAL and 0.1 mg/kg NIC, 0.4 mg/kg of NIC increased locomotor activity in both male and female rats on days 1 and 4. However, only males exhibited NIC SST on day 4 relative to day 1 of exposure (within groups) and between groups on the test for sensitization two weeks later (Figure 2a, b). The locomotor response in females was greater than that of the male rats throughout testing, but it did not increase from day 1 to day 4 nor was there a significant group difference on the test for sensitization. Analysis of the time course data from the 2-hour sensitization test showed that the lack of SST in females was not due to a ceiling effect (Figure 2a, b). Thus, F344/BN F1 rats are responsive to NIC exposure, which has a long-lasting effect that leads to male specific NIC SST.

**Figure 2.**
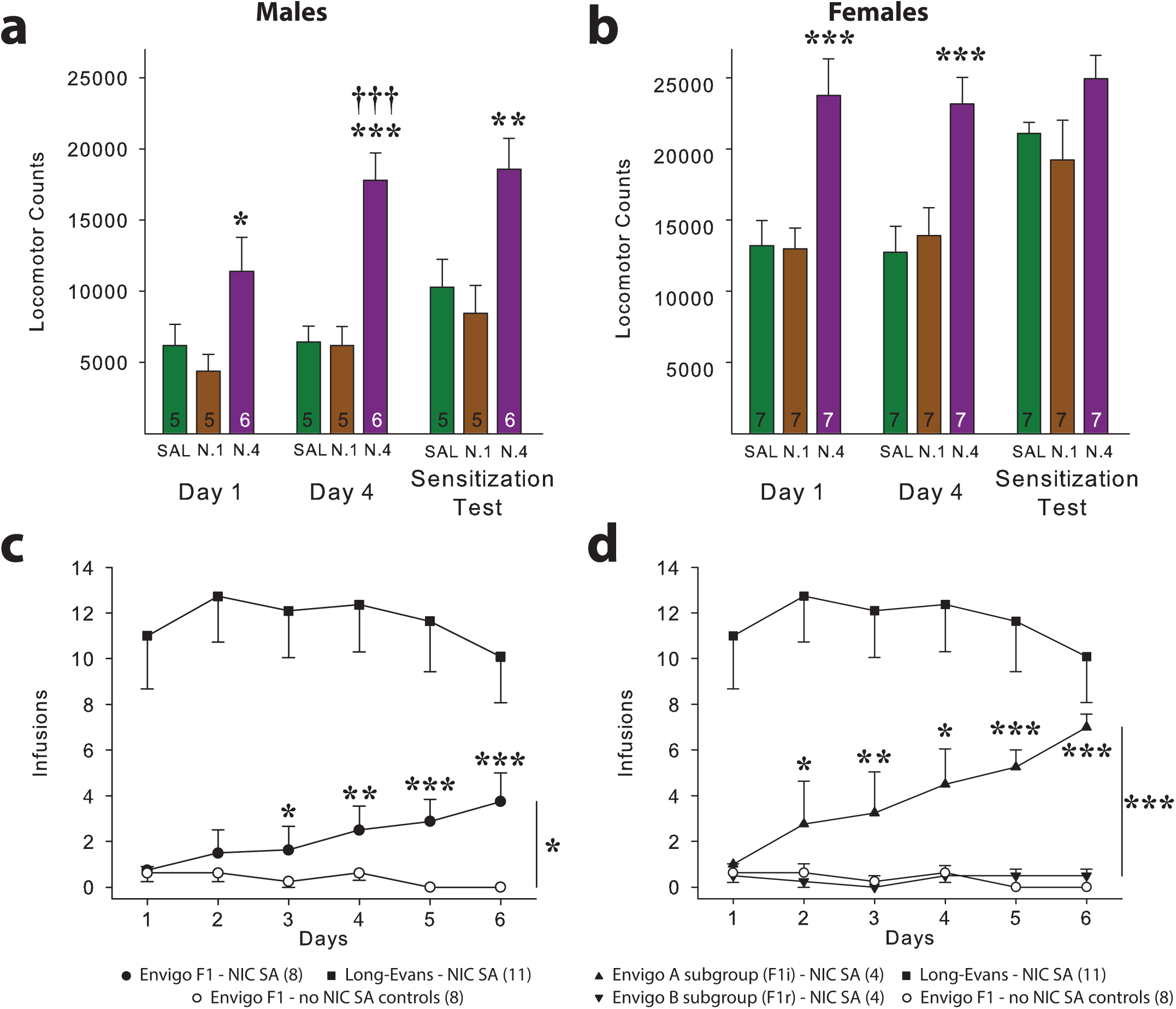
NIC sensitization (SST) and self-administration (SA) behavioral tests. (**a, b**) F1 males (n=5-6/group) and F1 females (n=7/group) were administered NIC (0.1 [N.1] or 0.4 [N.4] mg/kg; base, IP) or SAL daily for 4 days and tested for SST 2 weeks later. Data are mean (+SEM) of 2-hr total locomotor counts obtained on days 1 and 4, and on the test for SST when all rats were administered NIC (0.4 mg/kg). (**a**) Males showed a dose-dependent increase in NIC-induced locomotion and NIC SST. ANOVA of the day 1 and day 4 results in the males revealed significant effects of group [F_2,13_=10.43 (*p*<0.01)], days [F_1,13_=15.55 (*p*<0.01)], and a significant group X days interaction [F_2,13_=7.11 (*p*<0.01)]. ANOVA of the test for sensitization results in these rats showed a significant group effect [F_2,13_=7.25 (*p*<0.01)]. The denoted *p*-values were from post hoc Scheffé comparisons: **\***, *p*<0.05, **\*\***, *p*<0.01, **\*\*\***, *p*<0.001, N.4 vs two other groups at indicated days. **†††**, *p*<0.001, day 4 vs day 1 in N.4. (**b**) Females showed a dose-dependent increase in NIC-induced locomotion but did not exhibit NIC SST. ANOVA yielded only a significant effect of groups [F_2,18_=17.74 (*p*<0.001)] for the exposure day 1 and 4 results in females. The denoted *p*-values were from post hoc Scheffé comparisons: **\*\*\***, *p*<0.001, N.4 vs two other groups at indicated days. (**c**) Envigo F1s with access to NIC as a group self-administered the drug significantly more than the non-catheterized no NIC self-administration Envigo controls but much less than outbred Long-Evans rats (from ^28^; illustrated here for comparison). ANOVA of the results obtained in the two Envigo groups with and without access to NIC revealed a significant effect of groups [F_1,14_=9.35 (*p*<0.05)] and a significant group X days interaction [F_5,70_=4.17 (*p*<0.01)], with post Scheffé comparisons showing a progressively increasing and significantly higher intake in the rats with access to NIC starting on day 3 of SA (*p*<0.05-0.001). (**d**) When the F1s with access to NIC were divided by type of reciprocal cross [F344 father/BN mother F1s (subgroup A) and F344 mother/ BN father F1s (subgroup B)] and the results reanalyzed, subgroup A F1 rats showed more inclined NIC SA that approached levels seen in the Long-Evans outbred rats by day 6. Subgroup B F1s showed little to no NIC SA throughout testing. The ANOVA conducted on these results (in subgroups A and B and the no NIC self-administration controls) revealed significant effects of groups [F_2,13_=20.35 (*p*<0.001)], days [F_5,65_=5.16 (*p*<0.001)], and a significant groups X days interaction [F_10,65_=6.51 (*p*<0.001)], with post hoc Scheffé comparisons showing a progressively increasing and significantly higher intake only in subgroup A relative to the other two groups starting on day 3: * *p*<0.05, ** *p*<0.01, *** *p*<0.001. Data in (**c**,**d**) are the group mean (±SEM) number of infusions rats self-administered.

Because only males show SST, we modeled SA only in male F1 rats. The opportunity to lever press for NIC (30 µg/kg/infusion, IV) were given on each of 6 daily sessions. Again, reciprocal crossed rats (F1i and F1r) were evenly distributed in each group. As shown in Figure 2c, the Envigo F1s as a group emitted more lever presses for NIC than the non-catheterized no NIC self-administration controls, but much less than outbred Long-Evans rats (from ^28^ for illustration). This is in line with the literature showing that BN rats do not self-administer drugs as well as outbred strains like the Long-Evans ^16, 24^. However, when the data were reanalyzed by dividing the F1s by the type of reciprocal cross (i.e., A and B subgroups; Figure 1a), we surprisingly found rats in subgroup A (F1i: F344 father/BN mother F1s) were more inclined to self-administer NIC while rats in subgroup B (F1r: F344 mother/ BN father F1s) were disinclined to self-administer NIC (Figure 2d). The NIC SA observed in the F1i rats of subgroup A approached levels seen in the Long-Evans rats while the F1r rats in subgroup B showed little to no lever pressing (Figure 2d). These results indicate that F344/BN F1 rats self-administer NIC with a strong parental effect.

### NIC SST transcriptomic analysis identifies the VTA as the most relevant brain region

Because only male F1 rats exhibited NIC-induced locomotor SST despite the females showing a greater locomotor response to NIC (Figure 2a, b), we analyzed the transcriptomic profile associated with NIC SST mainly in males. We also conducted the transcriptomic analyses in females as a comparison to infer the molecular mechanism of sex-specific NIC SST. To minimize the confounding transcriptomic effect of acute NIC, F1 rats were sacrificed two weeks after the last NIC/SAL exposure without administrating NIC before harvesting VTA, NAc core and shell tissues for RNA-seq. Principal component analysis (PCA) of the transcriptomic data showed clear brain region-specific clustering, with VTA samples more distinct from NAc core and shell (Figure 3a), consistent with preserved region-specific gene expression. Brain region identity was also confirmed by the expression profiles of a set of genes related to NIC function (e.g., nAChRs expressed in VTA; Extended Data Fig. 1). As expected, the RNA-seq samples were also well-separated by sex (Figure 3a, b).

**Figure 3.**
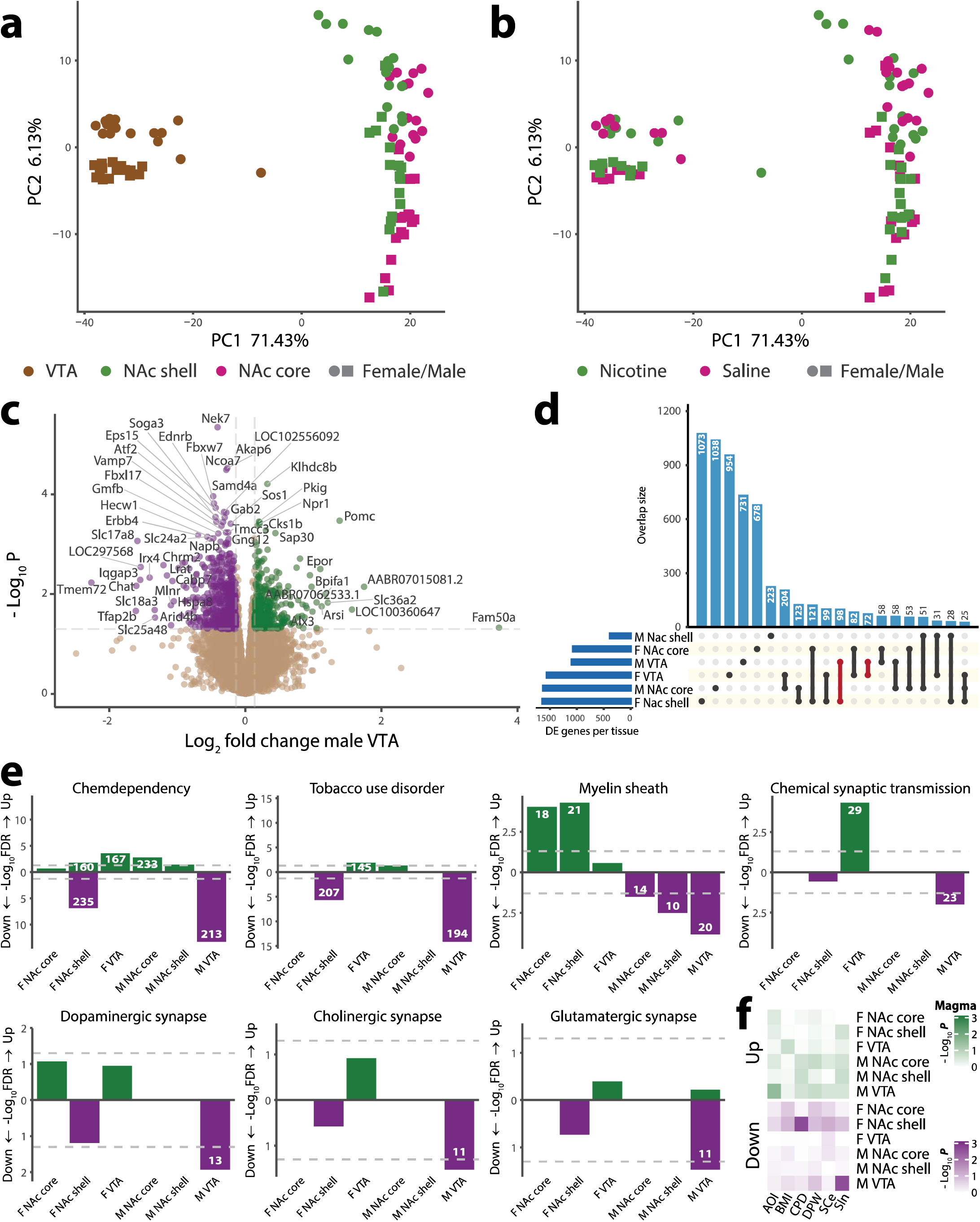
Transcriptomic analysis of NIC sensitization (SST). (**a**,**b**) Principal component analysis (PCA) of the top 500 differentially expressed (DE) genes in response to SST colored by (**a**) brain region, and by (**b**) NIC treatment; NAc – nucleus accumbens, VTA – ventral tegmental area. (**c**) Volcano plot of DE genes in the male VTA. (**d**) Upset plot of genes that are DE in different brain regions, with a nominal *p*<0.05. Overlaps with n≥25 are shown. (e) DAVID gene set enrichment analysis of DE genes in each brain region, examining GAD diseases and disease classes, OMIM diseases, KEGG pathways and GO terms. FDR-significant gene sets include the number of class genes (inset in the bar). (**f**) MAGMA enrichment analysis of genes harboring GWAS risk variants of five addiction phenotypes ^11^ among DE genes in each brain region. Gene interval – 100kb window around the gene. AOI – age of initiation, BMI – body mass index, CPD – cigarettes per day, DPW – drinks per week, SCe – smoking cessation, SIn – smoking initiation; Up – upregulated genes, Down – downregulated genes; M – male, F – female.

To determine the transcriptomic profiles associated with NIC SST, we identified differentially expressed (DE) genes in each brain region between NIC-treated and SAL groups. We found that the expression fold changes associated with NIC SST were overall relatively small (<2-fold) (Figure 3c, Extended Data Fig. 2), consistent with a polygenic nature of NIC addiction traits. With a relaxed statistical cut-off (*p*<0.05), we found 1629, 386, and 1097 DE genes out of 16,068 expressed genes in male NAc core, shell, and VTA (Supplementary Table 1). Females showed a comparable 1074 and 1562 DE genes in the NAc core and VTA respectively but more (1643) DE genes in NAc shell (Supplementary Table 2, Extended Data Fig. 2). The expression changes for some selected genes were independently confirmed by quantitative PCR (qPCR) (Extended Data Fig. 3). Overall, there was little overlap of DE genes between different brain regions and between sexes (Figure 3d and Extended Data Fig. 2), consistent with the notion of brain region-specific and sex-specific expression in transcriptomic PCA (Figure 3a).

**Table 1.**
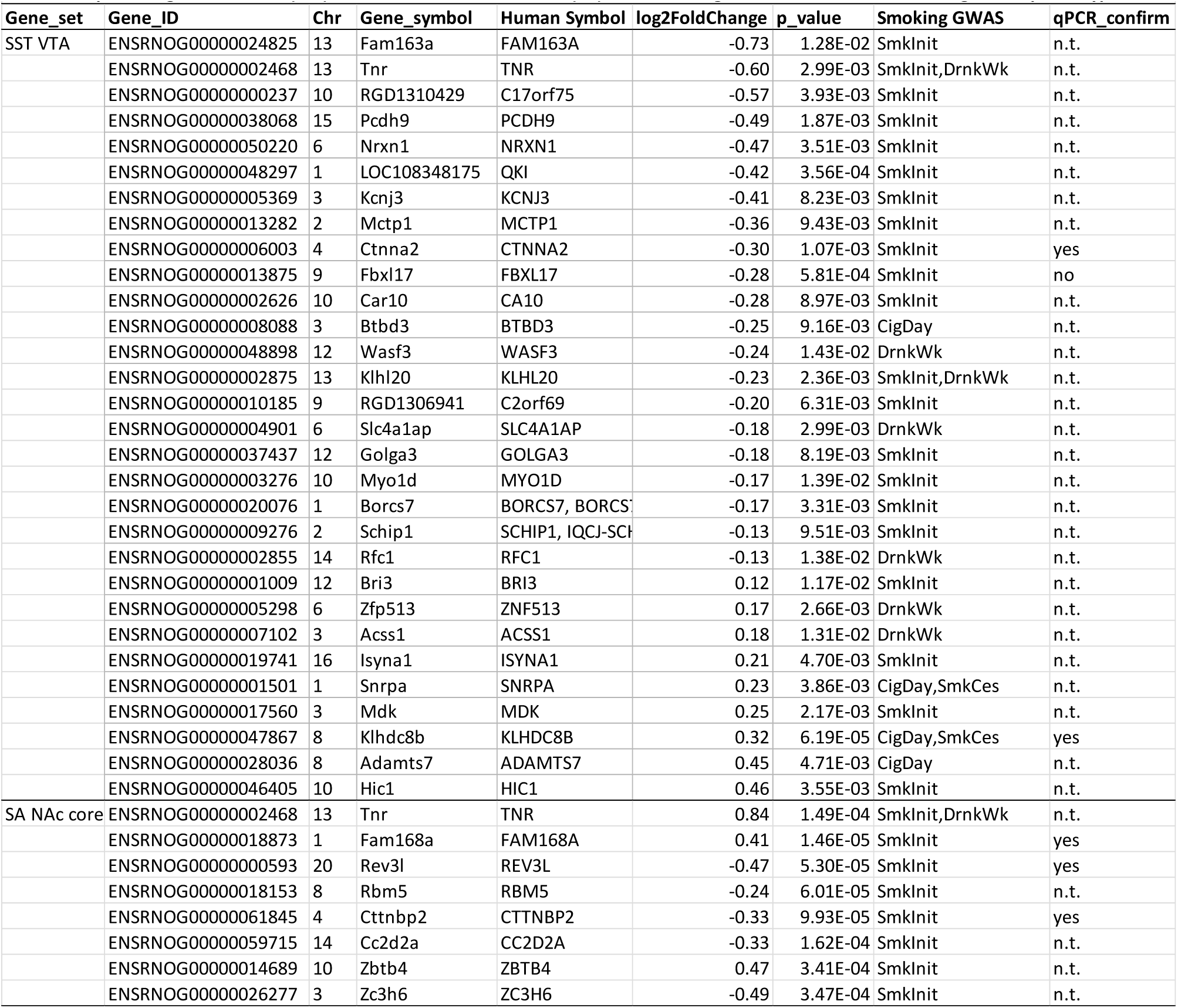
Top-ranking sensitization (SST) VTA and self-administration (SA) NAc core DE genes associated with smoking GWAS phenotypes.

We next assessed the functional properties of DE genes in each region to determine those that are most relevant to NIC SST. Gene set enrichment analysis was performed with human orthologs of each DE gene, using DAVID 6.8 ^25, 26^ to check for enrichment of GAD/OMIM disease classes, gene ontologies (GO), and KEGG pathways. We found that the down-regulated genes in the male VTA showed strong enrichment for genes related to tobacco use disorder and chemdependency, as well as NIC-related brain function, including myelin sheath, chemical synaptic transmission, postsynaptic density, dopaminergic/glutamatergic/cholinergic synapse, and nervous system development (Figure 3e, Extended Data Fig. 4). In contrast, up-regulated genes in the VTA showed no (for males) or a lesser (for females) enrichment of GO terms related to brain function (Figure 3e, Extended Data Fig. 4). Interestingly, although female F1 rats did not show SST (Figure 2b), significant enrichments of gene sets related to NIC addiction and brain function were observed in female NAc shell (Figure 3e), suggesting sex-specific lasting NIC effects in these brain regions.

To ascertain the genetic relevance of SST-associated DE genes to NIC addiction phenotypes in humans, we performed a MAGMA ^27^ gene-set analysis to test for enrichment of GWAS signals of tobacco and alcohol use disorders ^11^. MAGMA first maps GWAS risk loci to individual genes and then computes an enrichment of GWAS associations for a gene set. We found that down-regulated genes in the male VTA and female NAc shell showed enrichment (*p*<0.003) for SIn and CPD, respectively (Figure 3f). Out of the 100 top-ranking DE genes in the male VTA, ∼30% are associated with smoking GWAS phenotypes (with genome-wide significance, i.e., *p*<5×10^−8^), mainly SIn (Table 1); most of which are down-regulated genes with brain functions potentially related to NIC addiction. For instance, *Pcdh9* is critical for cognitive functions required for long-term social recognition ^28^; the knockdown of *LOC108348175* (*Qki*) impairs and enhances dendritic formation ^29^; and *CTNNA2* is a key regulator of the stability of synaptic contacts ^30^.

Taken together, these results demonstrate a predominant role of gene expression in the VTA region in mediating the long-lasting neural synaptic effects of NIC on male specific NIC SST. Furthermore, the transcriptional changes underlying NIC SST are relevant to human smoking phenotypes, mainly SIn.

### Sex-specific NIC SST may be partially explained by opposite transcriptional changes in the VTA

To understand the molecular basis of the observed sex difference of NIC SST (Figure 2a, b), we first examined the correlation of NIC-induced expression changes in each brain region between male and female F1 rats. We observed a negative correlation (R=-0.25, *p*<1.48×10^−218^) of global NIC-induced gene expression between males and females in the VTA (and NAc shell), but not in the NAc core (Extended Data Fig. 5a-c). A stronger negative correlation of expression changes was found for the subset of overlapping DE genes in male and female VTAs (R=-0.88, *p*<5.65×10^−39^; n=115, Supplementary Table 3, Figure 4a). On the contrary, no significant correlation of gene expression changes was found between male VTA and female NAc shell (Figure 4b), a brain region that also showed transcriptional relevance to long-lasting NIC effect in females (Figure 3e). The strong negative correlation of DE genes between male and female VTA was not due to their baseline (i.e., SAL-treated) expression in males and females, which showed very strong positive correlation (R=0.99, *p*<6.42×10^−145^) (Extended Data Fig. 5d). Therefore, the overlapping DE genes in VTA of male and female rats with opposing expression changes likely contribute to the sex difference of NIC SST.

**Figure 4.**
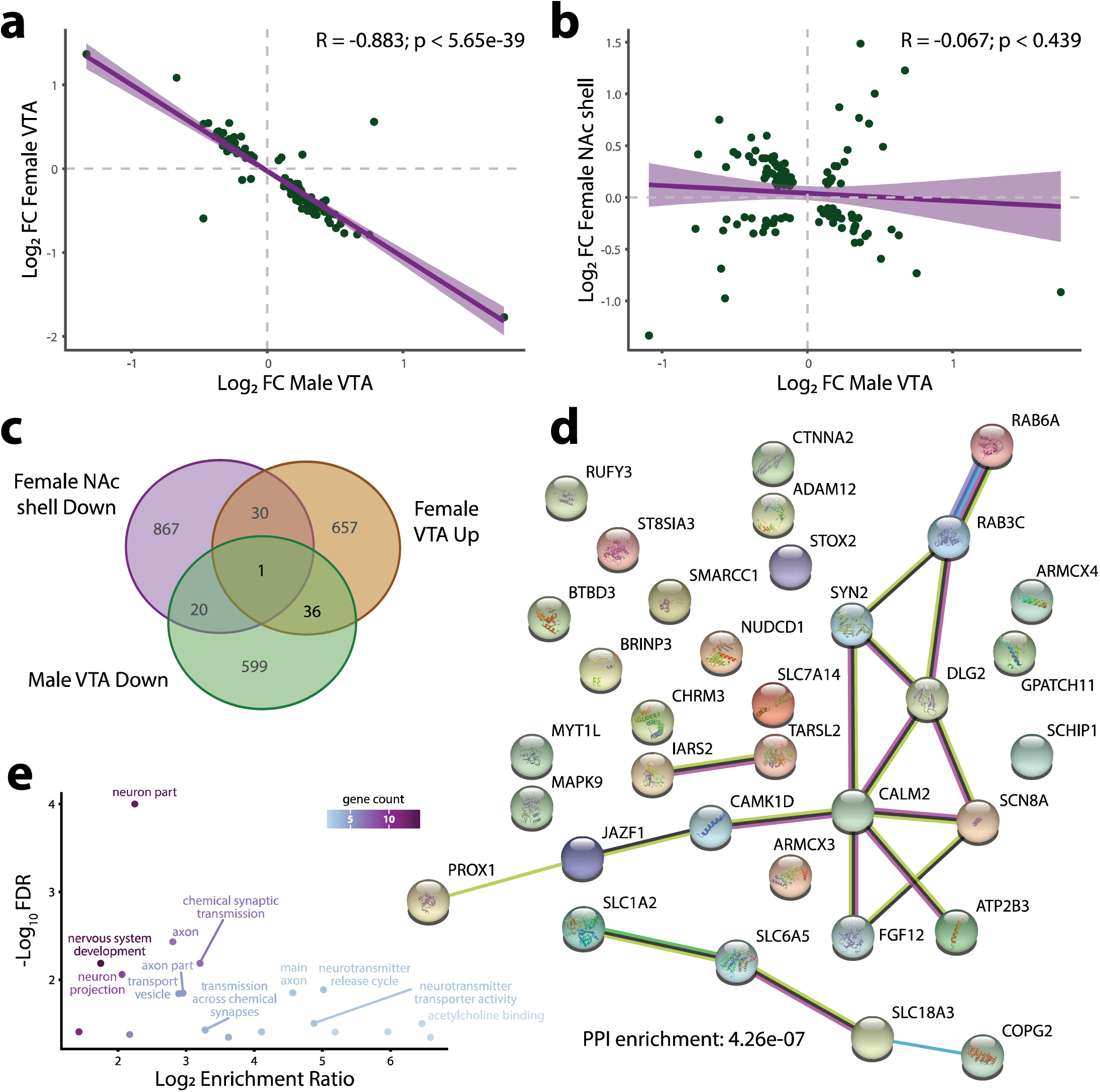
Comparison of sensitization (SST)-associated differential expression (DE) between males and females. (**a**) Fold change comparison of genes with nominally significant (*p*<0.05) DE in both ventral tegmental areas (VTAs) between males and females. (**b**) Fold change comparison of genes with nominally significant (*p*<0.05) DE in the male VTA and female NAc shell. (**c**) Venn diagram of nominally significant (*p*<0.05) DE genes in directions/regions enriched for addiction phenotypes. Note: male VTA downregulated, female VTA upregulated, and female NAc shell downregulated genes are most relevant to NIC addiction based on DAVID and MAGMA enrichment analyses (Figure 3, Extended Data Figs. 2, 5). (**d**) STRING analysis of human orthologs of 37 genes with opposite direction of DE (*p*<0.05) in male (down) and female (up) VTAs. Number of nodes: 34, number of edges: 18, average node degree: 1.06, avg. local clustering coefficient: 0.305, expected number of edges: 4, PPI enrichment *p<*4.3×10^−7^. (**e**) Ontological enrichments from the STRING analysis, colored by gene count in each term.

Given that male VTA downregulated and female VTA upregulated genes were enriched for GO terms related to NIC addiction and brain function (Figure 3e and Extended Data Fig. 4), it is conceivable that the small subset of 37 male VTA down/female VTA up genes (Figure 4c) with negative correlation of DE (Figure 4a) have a predominant effect on male specific NIC SST. In support of this hypothesis, we found that 5 out of 37 overlapping genes (13.5%) have nearby SNPs associated with smoking GWAS phenotypes, CPD, and/or SIn (Supplementary Table 3), representing 3.3-fold enrichment vs. 4.1% among all expressed genes (Fisher’s exact test, *p*<0.02). The hypothesis was also supported by our STRING ^31^ protein analysis of these 37 genes that showed a significant enrichment (18 edges vs. 4 expected, *p*<4.3×10^−7^) of protein-protein interaction (PPI) (Figure 4d), suggesting these genes are functionally more related to each other. It is noteworthy that the enriched PPI networks centered at *Slc18a3*, a vesicular acetylcholine transporter, and at *Camk1d (calcium/calmodulin-dependent protein kinase type 1D)* and *Calm2* (calmodulin-2) (Figure 4d). Further gene-set enrichment analysis ^32^ showed strong enrichment of GO-terms related to neuronal function (Figure 4e).

The importance of these 37 Mvta/Fvta negatively correlated genes (Figure 4c) in sex-specific SST was further supported by an enrichment of chromosome X (ChrX) genes. We found that 4 (*LOC100910130, Atp2b3, Armcx4, Armcx3*) out of the 37 negatively correlated genes (10.8%) are in ChrX (Supplementary Table 3); compared to 540 ChrX genes among all expressed genes (3.4%), this represents a 3.2-fold enrichment of ChrX genes (Fisher’s exact test, *p*<0.03). Interestingly, *Atp2b3*—one of the ChrX genes that is also part of the enriched PPI network involved in synaptic calcium signaling (Figure 4d)—encodes a calcium pump that has been suggested to be responsible for sex-specific pain responsiveness through interaction with estrenog recetpeor alpha(ERα) in mice ^33, 34^. Altogether, these results suggest the NIC-induced opposing transcriptional changes of a small set of genes that are enriched for ChrX genes in male and female VTA may mechanistically contribute to the observed male-specific NIC SST.

### NIC SA transcriptomic analysis identifies NAc core and VTA as relevant brain regions

Previous transcriptomic studies of NIC SA in rodents often use strains with different genetic backgrounds that may confound the results ^35, 36^, or have an experimental setting that assay the acute effect of NIC exposure rather than the molecular changes that may lead to SA ^37^. Leveraging our unexpected observation that only the subgroup A (F1i, with F344 as paternal strain) of the reciprocally crossed male F344/BN F1 rats (Figure 1a) showed inclination for SA (Figure 2d), we analyzed SA-associated DE genes by directly comparing the transcriptomes of the A vs. B subgroups in three brain regions (NAc core, shell and VTA). To mitigate the possible confounding effect of NIC exposure, we ensured that all rats received 10 infusions of NIC per session by providing a sufficient number of passively administered priming infusions (see methods), and harvested brain tissues 3 days following the last SA session. Similar to the SST transcriptomic analysis (Figure 3a, b), PCA of SA transcriptomes showed clear separation of three brain regions with the VTA more separated from the other two (Figure 5a), but no clear separation between the A and B subgroups (Figure 5b). Out of 16,063 expressed genes in the 3 brain regions, we found 1534, 387, and 1070 DE genes (*p*<0.05) in NAc core, shell and VTA, respectively (Supplementary Table 4). Most DE genes showed small to modest fold-changes of expression (<2-fold) (Figure 5c; Extended Data Fig. 6a), suggesting small effects of many genes on SA behavioral trait. The expression changes of three selected genes (*Rev3l, Fam168a*, and *Cttnbp2*) were all confirmed by independent qPCR (Extended Data Fig. 3b, c). In addition, we found very few overlapping DE genes between brain regions (Figure 5d; Extended Data Fig. 6b), indicating robust region-specific effect.

**Figure 5.**
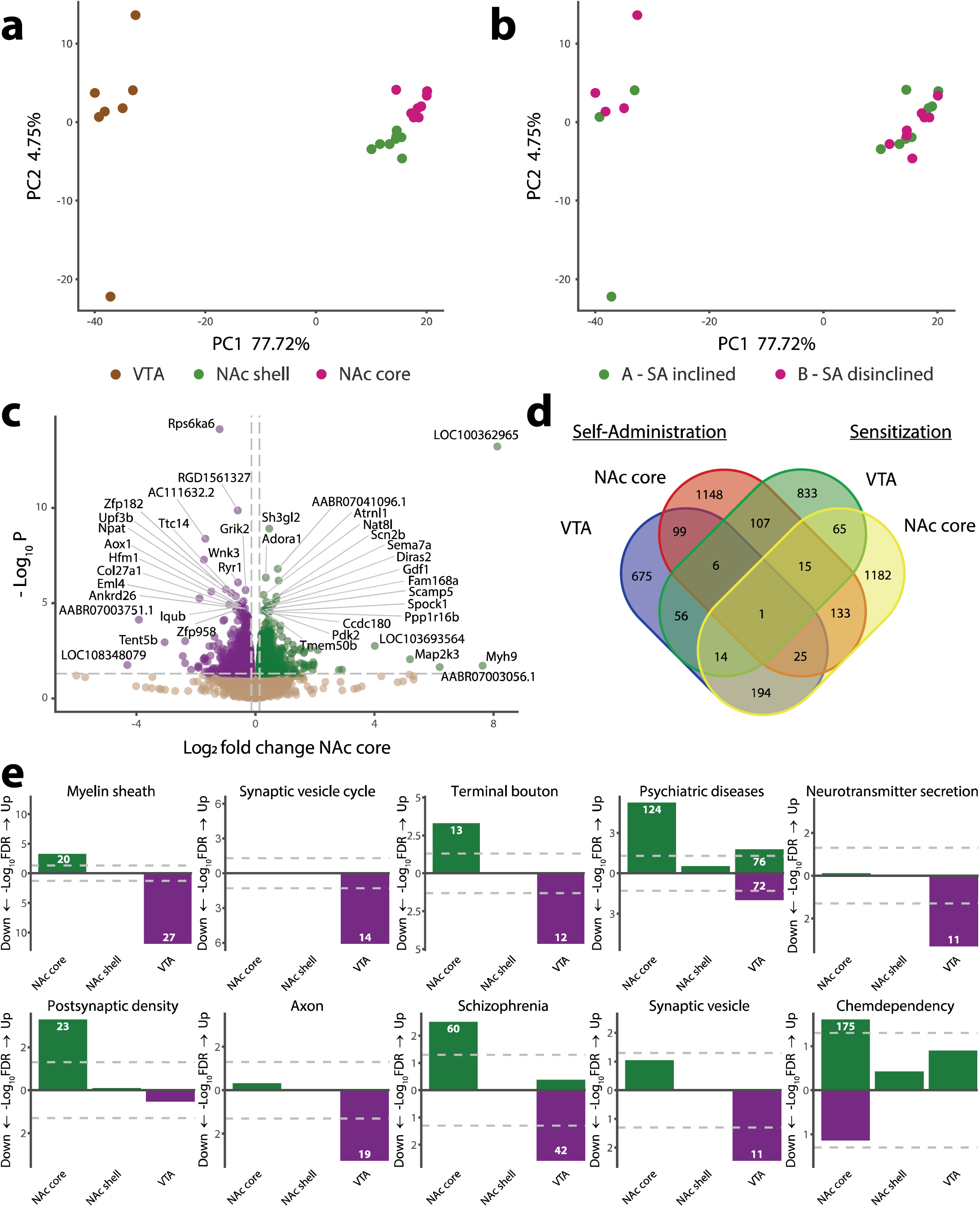
Transcriptomic analysis of NIC self-administration (SA). (**a**,**b**) PCA analysis of the top 500 differentially expressed (DE) genes in response to SA colored by (**a**) brain region and by (**b**) F1 cross subgroup (which experimentally was also the SA inclined/disinclined distinction); NAc – nucleus accumbens, VTA – ventral tegmental area. (**c**) Volcano plot of 15,443 DE genes in the male NAc core. (**d**) Venn diagram of the DE (*p*<0.05) genes in different brain regions. NAc core and VTA of SA experiments are compared to NAc core and VTA of SST experiments. **(e)** DAVID gene set enrichment analysis of DE genes in each brain region, examining GAD diseases and disease classes, OMIM diseases, KEGG pathways, and GO terms. FDR-significant gene sets include the number of genes (inset in the bar). Shown on the y-axis are enrichment significance (-Log_10_ FDR), Up – upregulated genes, Down – downregulated genes.

To identify the specific brain regions where DE gene sets are most relevant to NIC SA, we performed gene set enrichment analysis ^25, 26^ using human orthologs of DE genes. We found that both the NAc core and the VTA were biologically relevant (Figure 5e). Specifically, the upregulated genes in NAc core showed strongest enrichments for psychiatric diseases, chemdependency, and GO-terms such as neurotransmitter secretion, postsynaptic density, myelin sheath, and KEGG pathways including synaptic vesicle cycle (Figure 5e, Extended Data Fig. 7). In the VTA, downregulated genes were also strongly enriched for psychiatric diseases, and GO-terms related to neuronal function such as myelin sheaths, axon and neuron projection, neurotransmitter release, and synaptic vesicle (Figure 5e, Extended Data Fig. 7). MAGMA ^27^ gene-set enrichment analysis of NIC GWAS phenotypes (as shown in Figure 3f) did not produce conclusive results; however, VTA-down (downregulated) showed nominally significant enrichment (*p*<0.008) of AOI (not shown).

We next examined whether individual DE genes associated with NIC SA can help infer a likely risk gene at a smoking GWAS risk locus. Among the 100 top-ranking DE genes in the NAc core, 8 are associated with GWAS risk loci of SIn (Table 1). *Tenascin R* (*TNR*), which was the most upregulated gene in the NAc core (∼1.8-fold increase in A vs B subgroup), encodes an extracellular matrix glycoprotein that functions in neural cell adhesion and neurite outgrowth, regulates the balance of excitatory and inhibitory synapses ^38^, and is also associated with attention deficit hyperactivity disorder ^39^. The putative GWAS risk genes also included some downregulated genes in NAc core (Table 1). For instance, *Rev3l* has missense SNPs that are strongly associated with SIn (*p*<4.5×10^−29^) ^11^, and mutated *Rev3l* alters motoneuron migration or proliferation ^40^. *Cttnbp2*, another downregulated gene that modulates the density of neural dendritic spines ^41^, is also strongly associated with SIn (*p*<4.8×10^−22^) ^40^. Taken together, these results suggest that transcriptional changes in the NAc core and VTA during SA are relevant to NIC addiction and SIn in humans, and the identified DE genes may help infer the putative risk genes of smoking.

### NIC SA and SST are molecularly linked processes involving neurogenesis and myelin sheath

SA is a more commonly used model than SST for studying NIC addiction in rodents ^42^. It is largely unknown whether these two processes are mechanistically connected. Transcriptomic profiling of NIC SST and SA with the same set of male F1 rats enabled us to examine molecular links between the two processes. Comparing the DE genes in SST-relevant (VTA) and SA-relevant brain regions (NAc and VTA) showed a very small proportion of overlapping genes, even within the same brain regions (Figure 5d). However, for the overlapping genes between the most relevant regions for SST (VTA) and SA (NAc core), we surprisingly identified a significant negative correlation of gene DE associated with SST and SA (R=-0.50, *p*<2.24×10^−9^) (Figure 6a). The observed negative correlation of DE genes associated with SST and SA was largely driven by SA NAc core upregulated and SST VTA downregulated genes (Figure 6c and Supplementary Table 5), which are the most NIC addiction-relevant gene categories for each process. These negatively correlated genes accounting for 75% of the overlapping DE genes between SA NAc core and SST VTA (2.8-fold enrichment, Fisher’s exact test *p*<2.2×10^−16^) (Supplementary Table 5). For VTA, although we did not find a significant correlation of the DE genes between SST and SA (Figure 6b, Supplementary Table 6), we observed a strong positive correlation (R=0.66 and *p*<1.2×10^−10^) when a single extreme outlier gene (*Tmem72*) was removed. Of these 77 overlapping genes, 27 (35%) were downregulated in both VTAs. These observations suggest a possible molecular link between SST and SA in regions that are transcriptionally relevant to NIC addiction.

**Figure 6.**
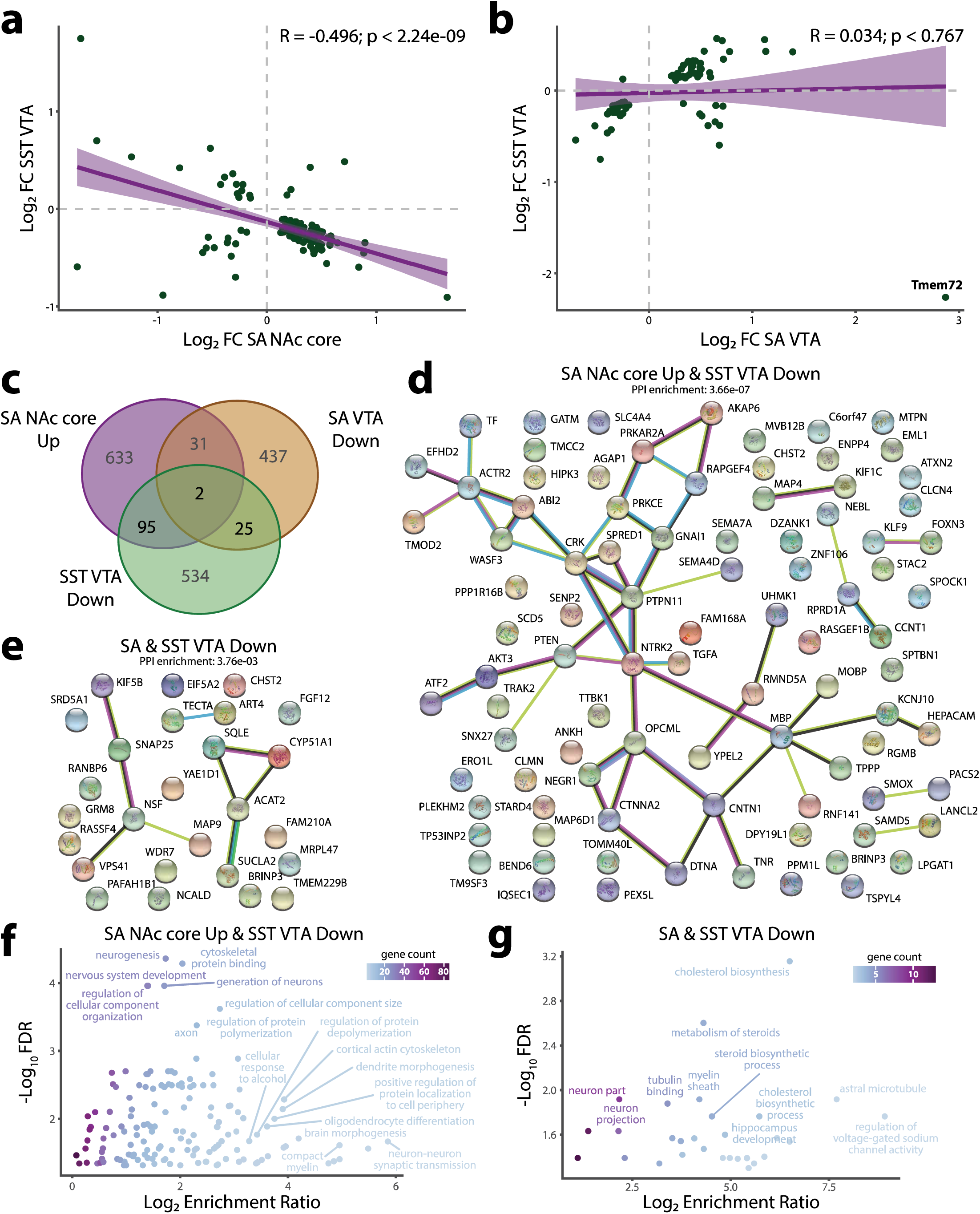
Comparison of differential expression (DE) of genes in regions enriched for addiction phenotypes between NIC sensitization (SST) and self-administration (SA). (**a**) Fold change (FC) comparison of genes with nominally significant (*p*<0.05) DE in the SST ventral tegmental area (VTA) and the SA NAc core (all from male rats). (**b**) FC comparison of genes with nominally significant (*p*<0.05) DE in the SST male VTA and the SA VTA. Note: excluding *Tmem72* (labeled), an expression outlier, gave a significantly positive correlation (R=0.66 and *p*<1.2×10^−10^). (**c**) Venn diagram of nominally significant (*p*<0.05) DE genes in directions/regions enriched for addiction phenotypes. Note: SST male VTA and SA VTA downregulated, as well as SA NAc core upregulated genes are most relevant to NIC addiction based on DAVID enrichment analyses (Figures 3, 5, Extended Data Fig. 6). (**d**) STRING analysis of the human orthologs of 97 genes with opposite DE direction in SA VTA (downregulated) and SA NAc core (upregulated). Number of nodes: 91, number of edges: 51, average node degree: 1.12, avg. local clustering coefficient: 0.333, expected number of edges: 23, PPI enrichment *p*<3.7×10^−7^. (**e**) STRING analysis of the human orthologs of 27 genes with concordant DE direction (downregulated) in SA VTA and SST male VTA. Number of nodes: 26, number of edges: 9, average node degree: 0.692, avg. local clustering coefficient: 0.321, expected number of edges: 3, PPI enrichment *p*<3.8×10^−3^. (**f**,**g**) Ontological enrichments from the STRING analysis in (**e**) and (**d**), respectively, colored by gene count in each term.

We next assessed the functional properties of the set of 97 overlapping SA NAc core-up (upregulated) and SST VTA-down genes, and the set of 27 overlapping SA VTA-down and SST VTA-down genes (Figure 6c). We reasoned that if these genes are important for both processes, they would likely form a PPI network relevant to NIC addiction. We thus carried out a STRING ^31^ protein analysis for the 2 sets of overlapping genes, and found significant enrichment in both for PPI network interactions (Figure 6d, e). Gene set enrichment analysis ^32^ further showed that both sets of overlapping genes are significantly enriched for GO-terms related to neurogenesis and myelin sheath (Figure 6f, g, Supplementary Tables 7, 8). Notably, myelin sheath was a GO term strongly enriched in genes associated with both SST and SA, in particular for male VTA-down gene (Figure 3e, 5e). We further evaluated the relevance of the 97 overlapping genes (SA NAc core-up and SST VTA-down) to human smoking phenotypes. We found that 10 such genes (*Tnr, LOC108348175, Ctnna2, Negr1, Wasf3,Tf, Fam168a, Prkar2a, Ptpn11*, and *Atxn*2) were within loci associated with different smoking GWAS phenotypes (mostly SIn); compared to that only 1 out of the 34 other overlapping DE genes was within smoking GWAS risk locus, this represents a 3.5-fold increase of possible smoking GWAS risk genes (Supplementary Table 5). Of the 27 SST VTA-down/SA VTA-down overlapping DE genes, although only one gene, *N-ethylmaleimide sensitive factor (Nsf)*, was associated with smoking GWAS risk, it is a strong risk gene for both cocaine dependence and locomotor SST ^43- 45^ as well as for alcohol use disorder (AUD) ^46^ (Supplementary Table 6) and was included in several enriched synaptic vesicle GO terms (Supplementary Table 7). The 27 genes also showed a strong enrichment for myelin sheath, neuron astral microtubules, and steroid/cholesterol biosynthesis—activities associated with myelination (Figure 6g). These results suggest NIC SST and SA may share gene pathways that involve neurogenesis and myelination, despite being two largely independent processes both relevant to NIC addiction.

### Parental effect of SA is associated with allelic imbalance of expression (AIE)

To ascertain the molecular mechanism of our observed parental effect of SA in F344/BN F1 male rats (i.e., only subgroup A showed inclined SA), we first examined the role of ChrX genes. We found that the DE genes associated with SA showed robust enrichment of ChrX genes (FDR=0.0004) in NAc core but not in VTA, two regions transcriptomically relevant to SA. In the NAc core region, out of the 100 most associated DE genes, there were 9 ChrX genes, representing a 2.9-fold enrichment (Fisher’s exact test, *p*<0.005) (Supplementary Table 4). However, given that most of the SA-associated ChrX genes (52/67) showed a reduced expression in SA-inclined subgroup A while the upregulated genes in the NAc core were associated with SA, ChrX genes unlikely play a major role in the parental effect of SA.

We hypothesized that the inclined SA of subgroup A was due to the dominant paternal (F344 strain) allelic effect (Figure 1a) on SA-relevant genes, manifesting as differential AIE between the A and B subgroups. Leveraging the heterozygous genetic background of F1 rats (i.e., informative for allele-specific analysis), we analyzed transcriptome-wide AIE in the A and B subgroups to identify genes that showed differential AIE. Using the transcribed heterozygous SNPs in F1 rats as proxies for regulatory variants, we compared the RNA-seq counts (reads) of the two parent alleles at a SNP site to identify AIE (see methods) (Figure 7a) in each brain region. We found that among ∼8000 transcribed bi-allelic SNPs present in three or more samples, about 47-55% showed either A or B subgroup AIE (FDR<0.05, binomial test) in different brain regions (Extended Data Fig. 8a, Supplementary Tables 9, 10). To identify more biologically meaningful AIE differences between A and B subgroups, we performed a two-sample proportion test (FDR<0.05), applying an additional threshold of a minimum difference (>0.1) of the fraction of reference allele (i.e., the copy from BN) between the two subgroups. We identified 719, 748, and 637 SNPs that showed differential AIE between the A and B subgroups in NAc core, shell and VTA, respectively. Although the reference allelic fractions of these SNPs were correlated between the A and B subgroups within each brain region (Pearson’s ∼R0.24) (Figure 7b, Supplementary Table 9), more than 80% of the AIE SNPs (in about 75% of the genes) were region specific (Extended Data Fig. 8b, c), suggesting strong brain region-specific parental effect on gene expression.

**Figure 7.**
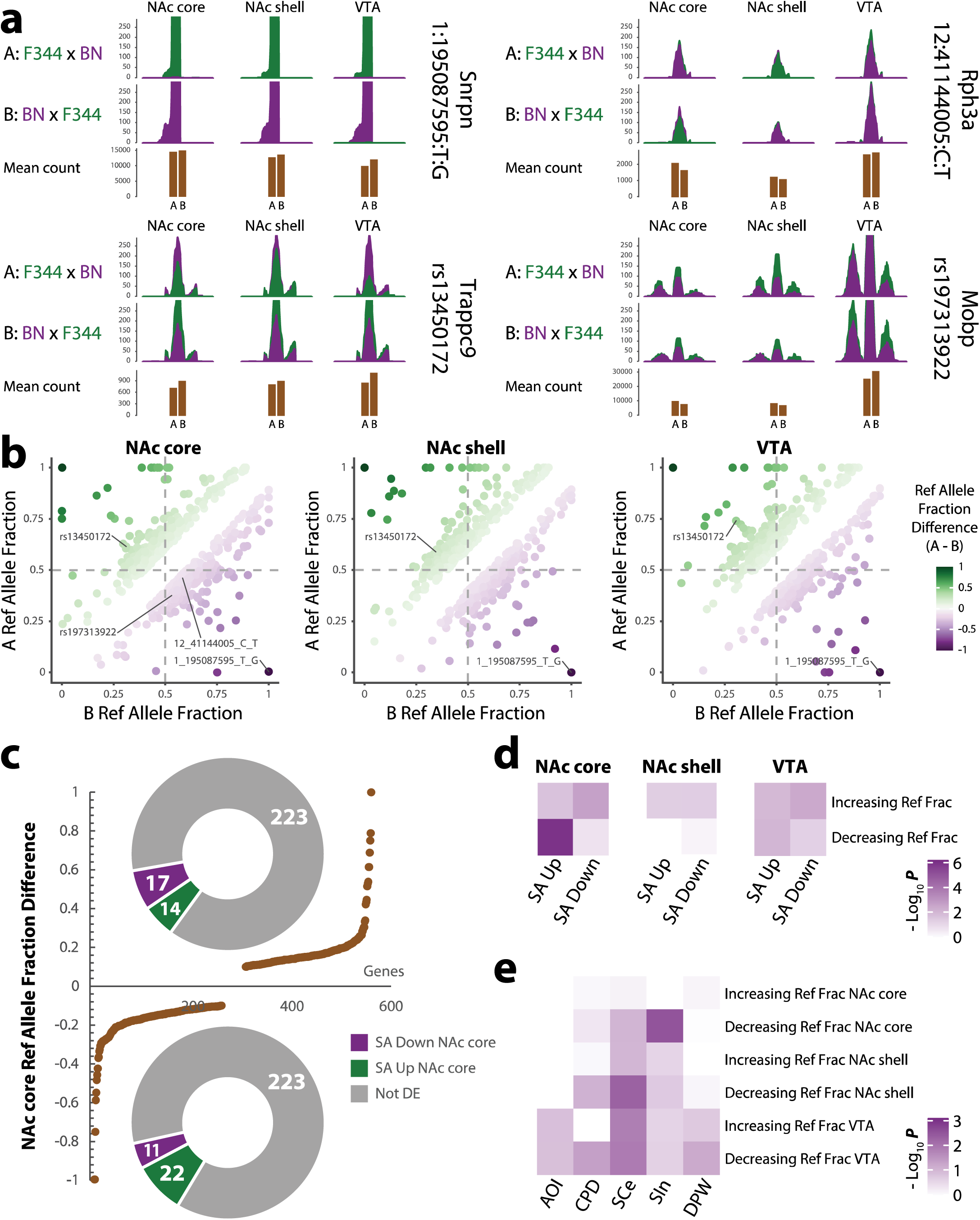
Allelic imbalance of expression (AIE) analysis of parental effect on self-administration (SA). (**a**) RNA-seq read pileup plots of example loci (transcribed SNPs) showing AIE in each brain region. The sequencing reads of the Fischer-344 (F344) allele are in green and those from the Brown Norway (BN) allele are in purple. Upper row depicts AIE of A subgroup (SA inclined), middle row depicts AIE of B subgroup (SA disinclined), and lower row depicts the normalized gene expression (mean RNA-seq read count) in A and B subgroups (shown in brown). (**b**) In each brain region (NAc core, shell and VTA; from left to right), correlation of the reference allele (BN copy) fraction of all AIE SNPs (binomial test, FDR<0.05; ref allele fraction difference>0.1) between the subgroup A and B rats. (**c**) Distribution of the reference allele (BN copy) fraction differences (A subgroup – B subgroup) of AIE genes in the NAc core (brown line). Donut plots show the number of genes in each DE category (upregulated – green; downregulated – purple; no change – gray) associated with AIE SNPs that showed either increased (above, >0.1) or decreased (below,<-0.1) reference allele fraction in subgroup A vs. B. (**d**) Heatmap showing the Fisher’s exact test enrichment of up- or downregulated genes in NIC SA among genes showing differential AIE (increasing or decreasing reference allele fraction; 2-sample proportion test, FDR<0.05) in subgroup A vs. B in each brain region. (**e**) Heatmap showing the Fisher’s exact test enrichment of genes harboring risk variants (r^2^>0.3 with GWAS index SNPs) of NIC addiction phenotypes ^11^ among AIE genes with significantly increasing or decreasing reference allele fraction (2-sample proportion test, FDR<0.05) in subgroup A vs. B in each brain region.

To identify genes associated with the parental effect on SA, we first examined those SNPs showing the largest difference of AIE (>70% allele fraction difference, A-B, i.e., near monoallelic in A or B) (Figure 7a, b). We found that such AIE SNPs with large parental effects were rare (n=12), and often showed consistent AIE patterns in all brain regions (Extended Data Fig. 8d, Supplementary Table 10). Many of these SNPs were found in genes or regions known to be imprinted in rodents or humans (Supplementary Table 10); for instance, *small nuclear ribonucleoprotein-associated polypeptide N (Snrpn)* ^47, 48^, *ubiquitin-protein ligase E3A (Ube3a)* ^49^, and *Trappc9* ^50^. AIE of *Snrpn* showed a strong bias toward the paternal allele of the SA-inclined subgroup B; correspondingly, *Snrpn* showed significant DE downregulation in the VTA (Figure 7a, b, Supplementary Table 10). *Trappc9* showed more mild AIE bias towards the maternal SA-disinclined allele of subgroup A across all tissues, with the strongest bias in the VTA, which had corresponding DE downregulation in both the NAc core and VTA (Figure 7a, b, Supplementary Table 10).

In the NAc core, we compared the genes showing a paternal AIE with those that were also upregulated. Overall, most genes showed small to modest AIE differences between the A and B subgroups and did not exhibit SA-associated differential expression (Figure 7c). However, we observed a strong enrichment (2.4-fold, Fisher’s xeact test *p*<2.1×10^−10^) of the SNPs with decreased AIE in the NAc core (Decreasing Ref Frac) among genes upregulated in NAc core of subgroup A (Figure 7c,d). Of the 22 overlapping genes (Figure 7c), 17 are associated with cognitive diseases, 5 are specifically associated with myelination (Fisher’s exact test, 30-fold, *p*<6.2×10^−7^), and 3 are within smoking GWAS risk loci (Fisher’s exact test, 4-fold, *p*<0.04) (Supplementary Table 11, Extended Data Fig. 9a). Further STRING network of this set of 22 genes (Extended Data Fig. 9b, c) identified myelin sheath as the top-ranking enriched GO that includes genes *Serinc5, Cntn2, Mobp*, and *Thy1* (Extended Data Fig. 9c, Supplementary Table 12).

Interestingly, genes with NAc core Decreasing Ref Frac SNPs also exhibited the strongest enrichment for GWAS association signals for SIn in MAGMA analysis (Fisher’s exact test, FDR<0.03) (Figure 7e). One such gene associated with SIn ^11^, *Rabphilin 3A (Rph3A)*, showed a decreased AIE while showing an increased expression in the NAc core of subgroup A rats (Figure 7a). *Rph3A* encodes a GluN2A subunit-binding partner that interacts with GluN2A and PSD-95, forming a complex that regulates N-methyl-D-aspartate receptor (NMDAR) stabilization at postsynaptic membranes and mediates strain-specific postsynaptic differences in mice ^51, 52^. Several of these 22 overlapping genes (Figure 7c), such as *Rapgef4, Enpp4*, and *Mobp*, also showed up in our STRING analysis of genes upregulated in the SA NAc core and downregulated in the SST VTA (Figure 6d). Taken together, these results suggest that the strain-specific AIE of autosomal genes in NAc core and VTA of F1 rats may mechanistically explain the observed parental effect of SA.

## Discussion

We modeled NIC addiction in F344/BN F1 rats, and surprisingly found that NIC SST was male-specific and there was a parental effect of NIC SA. By analyzing the transcriptomic signatures of NIC SST and SA in VTA, NAc core and shell regions, we found that the sex-specific NIC SST may be explained by genes with opposing NIC-induced expression changes in male and female VTAs, related to myelin sheath, synaptic transmission, and tobacco use disorders. For NIC SA, we found that both NAc core and VTA are transcriptionally relevant regions, involving genes related to neurogenesis, myelin sheath, postsynaptic density, and neurotransmission. Comparing the transcriptomic profiles of NIC SST and SA suggested a molecular link between these two independent processes related to NIC addiction. Our AIE analysis of NIC SA rats further demonstrated differential AIE between SA-inclined and SA-disinclined rats, which may mechanistically drive the differential gene expression associated with the parental or imprinting effect on SA. Finally, we found that DE genes associated with NIC SST and SA were enriched for GWAS associations with smoking phenotypes such as SIn and CPD, providing mechanistic and biological insights for many smoking GWAS risk loci.

The sex-specific effects on NIC SST have not been extensively studied and have yielded mixed results. For example, two previous reports show greater locomotor sensitization in female Sprague-Dawley rats with repeated intravenous nicotine administration ^53, 54^. By contrast, Pehrson and colleagues ^55^ report greater locomotor sensitization in male Sprague-Dawley rats with subcutaneous nicotine administration. This suggests that the route of nicotine administration may be an important variable in revealing sex differences in NIC SST. With F344/BN F1 rats we found that only male rats showed NIC SST, although female rats exhibited a strong locomotor response to NIC (Figure 2a, b). This is consistent with a previous report showing enhanced NIC locomotion in female relative to male Long Evans rats ^56^. Our transcriptomic analysis identified a strong negative correlation of NIC-induced expression changes between male and female VTAs (Figure 4a). Given that only the downregulated genes in male VTA are enriched for GO terms relevant to NIC addiction and brain function as well as GWAS risk variants of NIC SIn (Figure 4e, f), it is conceivable that these overlapping DE genes with opposite expression changes in VTA contribute to the sex-specific NIC SST. In support of this hypothesis, these overlapping genes were found enriched for ChrX genes related to neuronal function, e.g., *Atp2b3*, a gene that encodes a calcium pump implicated in sex-specific pain responsiveness in mice ^33, 34^. The enriched GO-term, myelin sheath, among these VTA genes (Figure 3e, Supplementary Table 13) also seems to be consistent with male-specific NIC SST: in rats, gestational exposure to NIC can produce sex-specific myelination effects during development ^57, 58^; in humans, sex-specific myelination has been reported to be correlated with differential brain development in boys and girls ^59, 60^, which may manifest into differential myelination-associated synaptic functionality ^61, 62^. Thus, the identified transcriptomic signature of male-specific NIC SST in rat may provide mechanistic insight into why men tend to use tobacco at higher rates than women ^63^ and smoking seems to activate male reward pathways more than in female ^64, 65^.

NIC SST and NIC SA have both been used to model NIC addiction in rodents. However, whether NIC SST and NIC SA are linked at the molecular level has not been explored. Our parallel transcriptomic analyses of NIC SST and SA in male F1 rats showed that VTA is an active site for both SST and SA, with downregulated genes enriched for chemical dependency and ontologies associated with myelin sheath and synaptic neurotransmission (Figure 3e, 5e). This is consistent with the notion that the VTA is an important region for the initiation of behavioral SST to psychostimulants ^15, 66-68^. In contrast to NIC SST, both VTA and NAc core showed transcriptional changes associated with NIC SA, with upregulated genes enriched for chemdependency, myelin sheath and postsynaptic density (Figure 5e). The molecular link between NIC SST and SA is further exemplified by the strong negative correlation of the expression changes of DE genes in VTA of NIC SST and in NAc core of NIC SA rats (Figure 6a), as well as the positive correlation of expression changes of DE genes in the VTAs of NIC SST and NIC SA rats (after removing extreme outlier *Tmem72*) (Figure 6b). Although the specific regulatory mechanism underlying the transcriptional correlation of DE genes within VTA or between VTA and NAc core for NIC SST and SA remains to be determined, we have found that the overlapping genes tend to form gene networks functionally related to neurogenesis and myelin sheath (Figure 6d-g), suggesting a strong molecular link between the two independent processes.

A parental effect on NIC SA has not been previously reported. The unexpected parental effect on NIC SA (Figure 2c, d) enabled us to identify the transcriptomic signatures of NIC SA by directly comparing the two subgroups of F1 rats, which was expected to give a more robust result because the two subgroups shared more similar experimental conditions than with the non-SA control group (Figure 2c,d). Despite the small sample size in each subgroup (n=4), we were able to identify DE genes associated with NIC SA in VTA and NAc core. To explore the molecular mechanism underlying the parental effect of NIC SA, we did not find evidence for a major role of ChrX genes; instead, we found that brain region-specific AIE SNPs in autosomal genes may explain the parental effect of NIC SA. Some of the genes showing strongest AIE differences between the A and B subgroups are known to be imprinted and likely contribute to the parental effect of NIC SA. For instance, *Snrpn*, a known paternally expressed gene ^47, 48^, and its bicistronic transcript partner *Snurf*, play an important role in adult neurogenesis ^69^ that is relevant to both NIC SST and SA (Figure 6f, g). *Snrpn* and *Snurf* are regulated via a paternal imprinting locus ^70^ and its dysregulation is associated with Praeder-Willi Syndrome ^71^ and inability to regenerate axons ^72^. Overall, our result fits with a model of paternal imprinting bias resulting in reduced expression in the VTA, and potentially implicates *Snrpn* in the downregulation of neurogenesis. *Trappc9* also demonstrates AIE, though seemingly in a maternal biased way, and is associated with downregulation in subgroup A (Figure 7a). Interestingly, a number of strong paternally biased AIE SNPs of *Trappc9* reside in an intron (Extended Data Fig. 10), whose corresponding human intronic region encompasses a paternally expressed long noncoding RNA (lncRNA) gene *PEG13* that is responsible for silencing the maternal transcription of *Trappc9* ^50, 73^. Further research into a potential *PEG13* rat ortholog would be warranted to decipher whether the subgroup A’s paternal bias toward *PEG13* results in altered expression and downregulation of *Trappc9*.

In aggregate, only AIE SNPs with reduced reference allele fraction in NAc core of subgroup A rats were enriched in genes upregulated in NAc core (Figure 7c, d), the most relevant gene set for NIC SA (Figure 5e). Interestingly though not necessarily connected, this AIE gene set with reduced reference allele fraction but upregulated in NAc core of subgroup A also shows an enrichment of GO terms of myelin sheath and axon (Extended Data Fig. 9c), and an enrichment of GWAS association signals for SIn (Figure 7e). Myelin sheath is also a GO term enriched in genes associated with SST and particularly with SA (Figure 5e, Supplementary Table 14). Notably, the *myelin-associated oligodendrocytic basic protein* (*Mobp*) gene, which showed paternally biased AIE and increased expression in the NAc core of subgroup A while with reduced expression in the VTA (Figure 7a), is a compelling target for further research to determine what role differential myelination may play in SST and/or SA. Overall, our AIE analysis further supports NAc core to be an essential region for SA, in particular its potential to be influenced by imprinting effects. Because the AIE SNPs here are proxies of transcription-regulatory variants, future study to identify putatively functional variants that may show differential allelic chromatin accessibility to transcriptional factors ^74, 75^ may shed light on the causal mechanism of parental effect on NIC SA-relevant upregulation of genes in NAc core.

For the first time, we have shown that the transcriptomic profiles of NIC SST and SA can inform the GWAS findings of human smoking phenotypes. Recent GWAS of human smoking phenotypes have identified hundreds of risk loci ^6-11^. However, it has been a challenge to distinguish the specific risk gene(s) from others that are equivalently associated with a phenotype at most of those risk loci. Our MAGMA analysis showed that the downregulated genes in male VTA in SST model were significantly enriched for GWAS risk SNPs of SIn (Figure 3f), with ∼30% of top-ranking DE genes of NIC SST in male VTA are associated with smoking phenotypes (Table 1). For NIC SA, although DE genes as a whole only showed nominally significant enrichment for GWAS risk SNPs of smoking AOI, the subset of AIE genes in NAc core were strongly enriched for GWAS risk SNPs of SIn (Figure 7e). These transcriptomic analyses thus not only orthogonally show the disease relevance of the rat NIC addiction models, but also highlight the value their transcriptomic signatures in helping dissect the biology underlying the GWAS findings of human smoking traits.

Given a possible gateway “drug” role of NIC ^4, 5^, our results may also have implications for understanding the genetic etiology and neurobiology of other drugs of addiction or substances of abuse. For instance, the expression changes of NIC SA-associated genes were strongly correlated (R^2^=0.22 for NAc core and R^2^=0.33 for VTA) with that of genes associated with cocaine SA ^76^ (Supplementary Table 4, Supplementary Fig. 1). Many upregulated genes in NAc core have neural functions such as neurogenesis (e.g., *Unc5b, Slc12a7, Sema5b, Adora1* and *Npas4)*. Notably, *Unc5b* and three other upregulated genes (*Tnrc6a,Tmx2, and Arid4a*) in NAc core of NIC SA rats (Supplementary Fig. 1a) are among those 26 GWAS risk genes for problematic alcohol use (PAU) or AUD ^77, 78^. It is also noteworthy that *CTNNA2*, a gene upregulated in the NAc core of SA rats and downregulated in the VTA of SST rats, is a key regulator of synaptic function ^30^ and associated with not only SIn ^11^, but also alcohol, heroin and methamphetamine dependence in Han Chinese ^79^.

In summary, our transcriptomic analyses suggested plausible mechanisms for the sex-specific NIC SST and the parental effect of NIC SA. Importantly, the DE genes associated with SST and SA are enriched for GWAS risk SNPs of smoking phenotypes. This study advanced our mechanistic understanding of NIC SST and SA, providing a valuable resource for narrowing down the biological relevant risk genes of smoking behaviors and other drugs of abuse disorders in humans.

---

**Extended Data Fig. 1.**
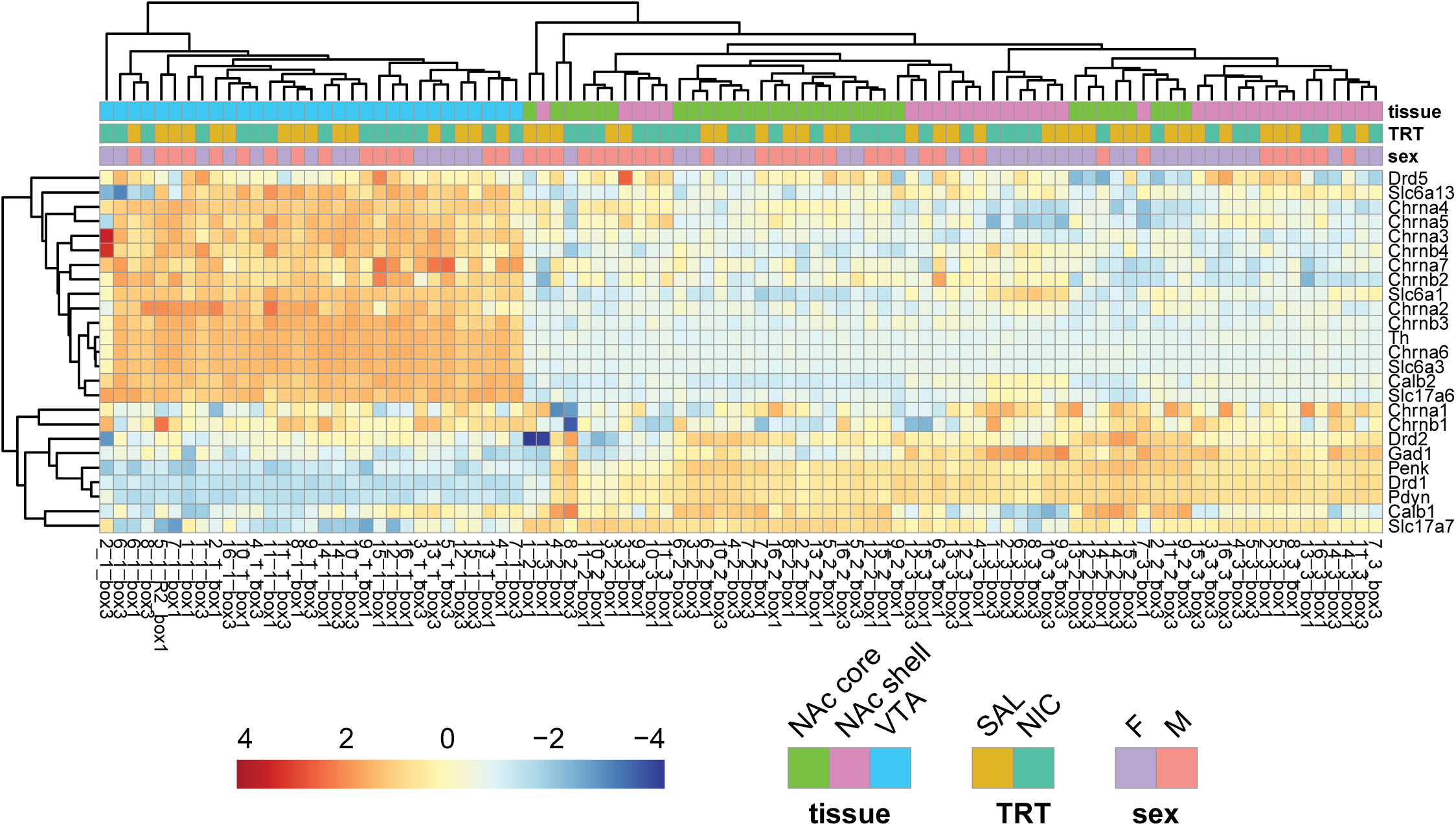
RNA-seq sample clustering based on a set of brain region-specific genes. Samples clustered by row-centered, variance stabilizing transform normalized values. Annotated by tissue (NAc core – nucleus accumbens core, NAc shell – nucleus accumbens shell, VTA – ventral tegmental area), treatment (TRT, NIC – nicotine injections, SAL – saline injections) and sex.

**Extended Data Fig. 2.**
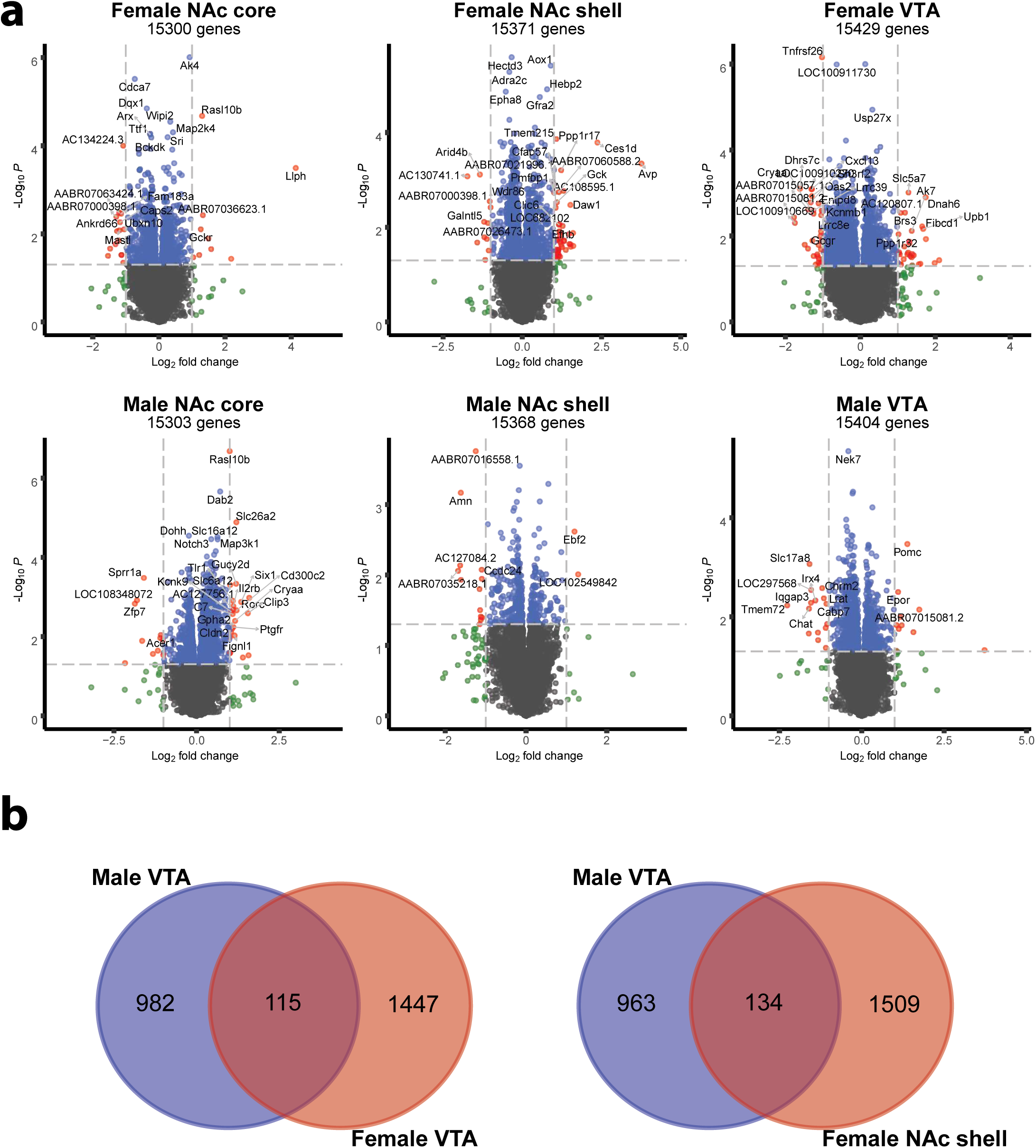
Differential expression (DE) of genes in each brain region of NIC-treated vs control F1 rats. (**a**) Volcano plots of DE genes in each brain tissue by sex. The horizontal dashed lines indicate 2-fold changes and the vertical dashed lines indicate the nominal significance *p*-value of 0.05. (**b**) Venn diagram showing the overlapping DE genes associated with NIC SST between males and females in regions transcriptionally relevant to tobacco use (determined from DAVID gene set enrichment analysis in Figure 3e).

**Extended Data Fig. 3.**
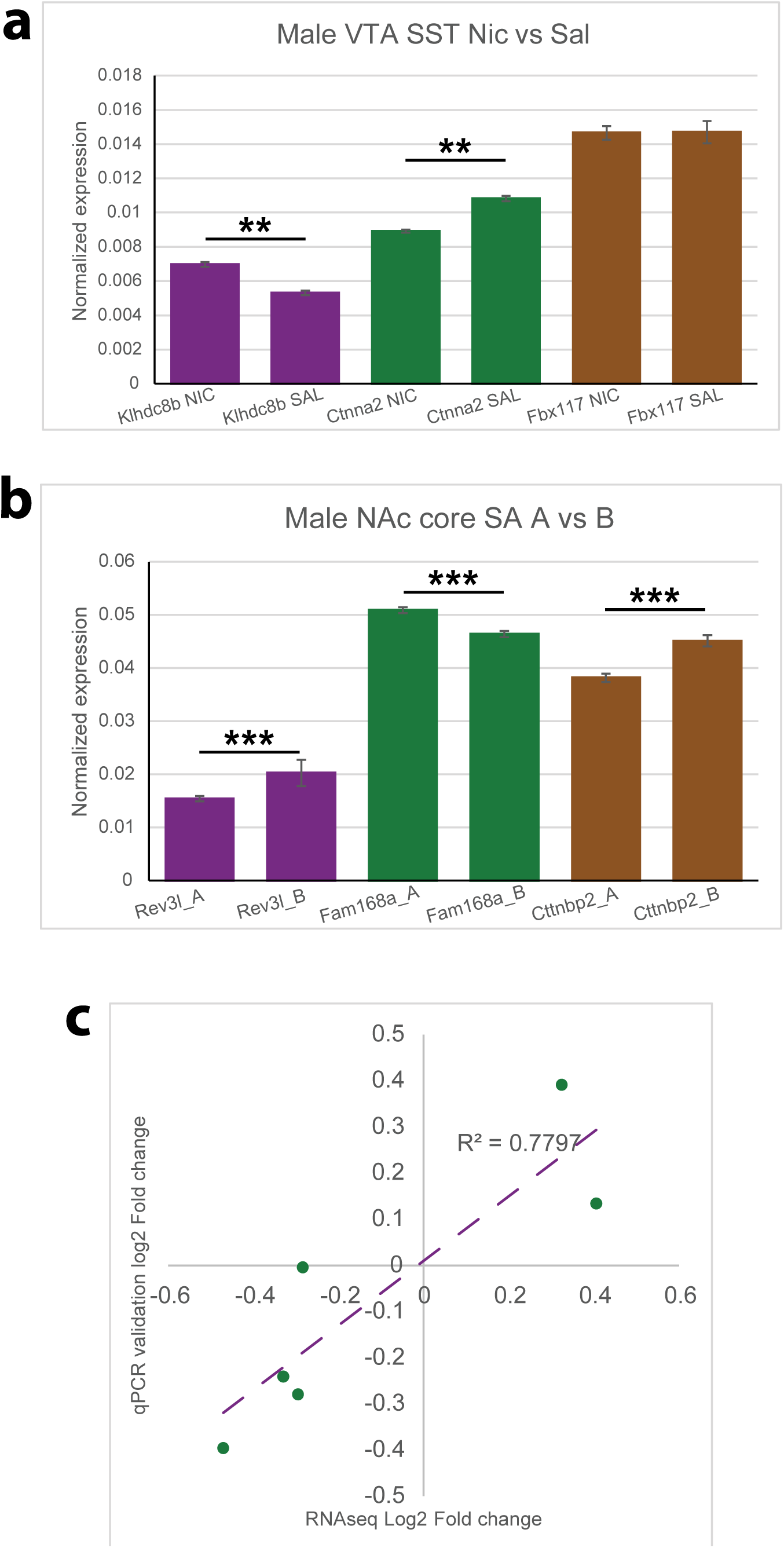
qPCR validation of the expression changes of 6 NIC SST-or SA-associated genes selected from RNA-seq DE analysis. (**a**) Two DE genes associated with SST (*Klhdc8b, Ctnna2*; ** *p*<0.03) and (**b**) three genes associated with SA (*Rev3l, Fam168a, Cttnbp2*; *** *p*<0.02) were confirmed by qPCR. The expression change of *Fbx117* in SST was not confirmed. Expression of each gene was normalized to *GAPDH*. Student’s *t*-test was used to test DE. Data are from 7 biological replicates (different animals) each with 3 technical replicates. Error bars, standard error of mean (SEM). **(c)** Strong correlation (R^2^=0.78) of gene expression fold changes of the selected genes between RNA-seq and qPCR validation.

**Extended Data Fig. 4.**
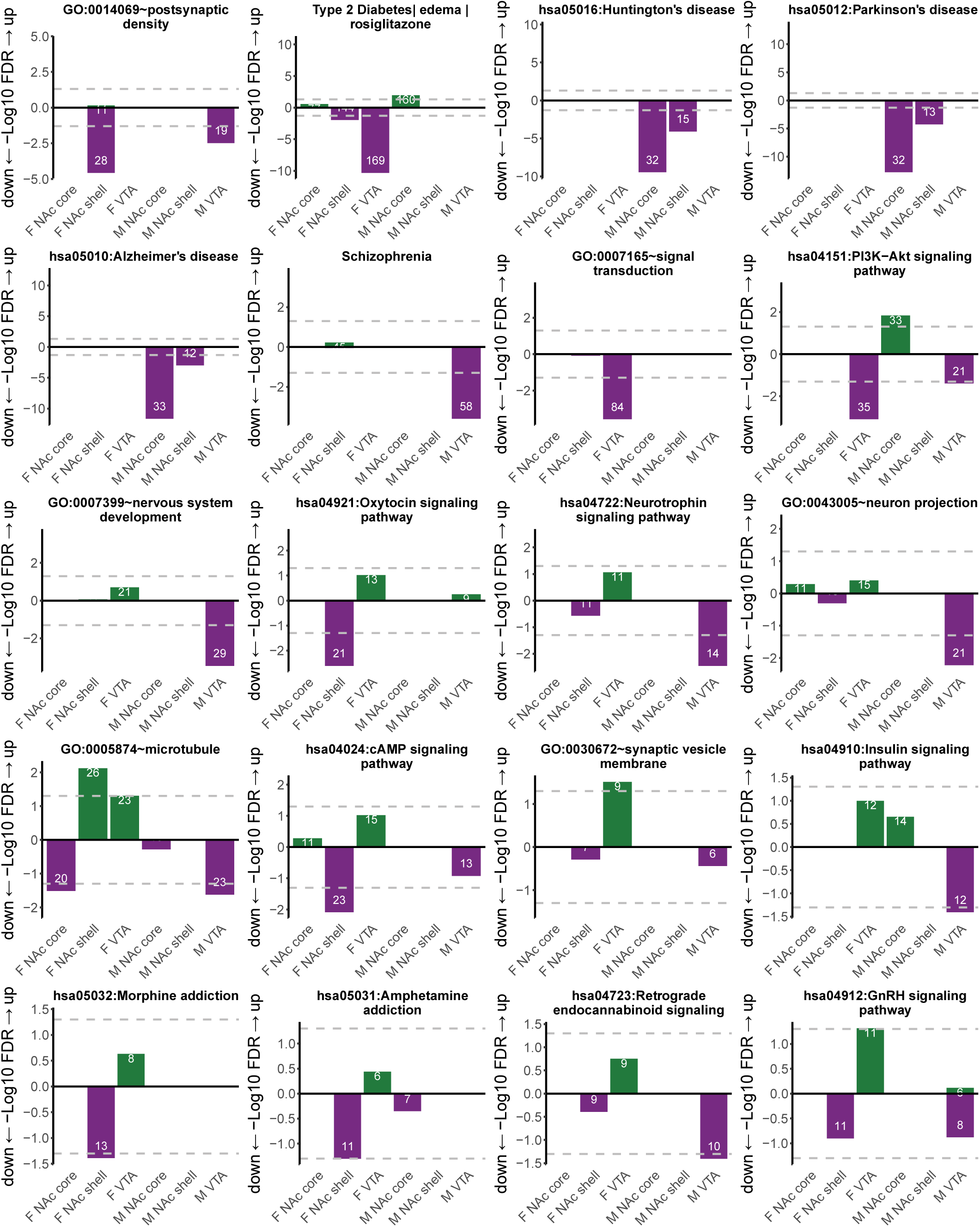
Extended DAVID gene set enrichment analysis of DE genes in each brain region for NIC sensitization. Listed are for GAD diseases and disease classes, OMIM diseases, KEGG pathways and GO terms. FDR-significant gene sets include the number of class genes (inset in the bar).

**Extended Data Fig. 5.**
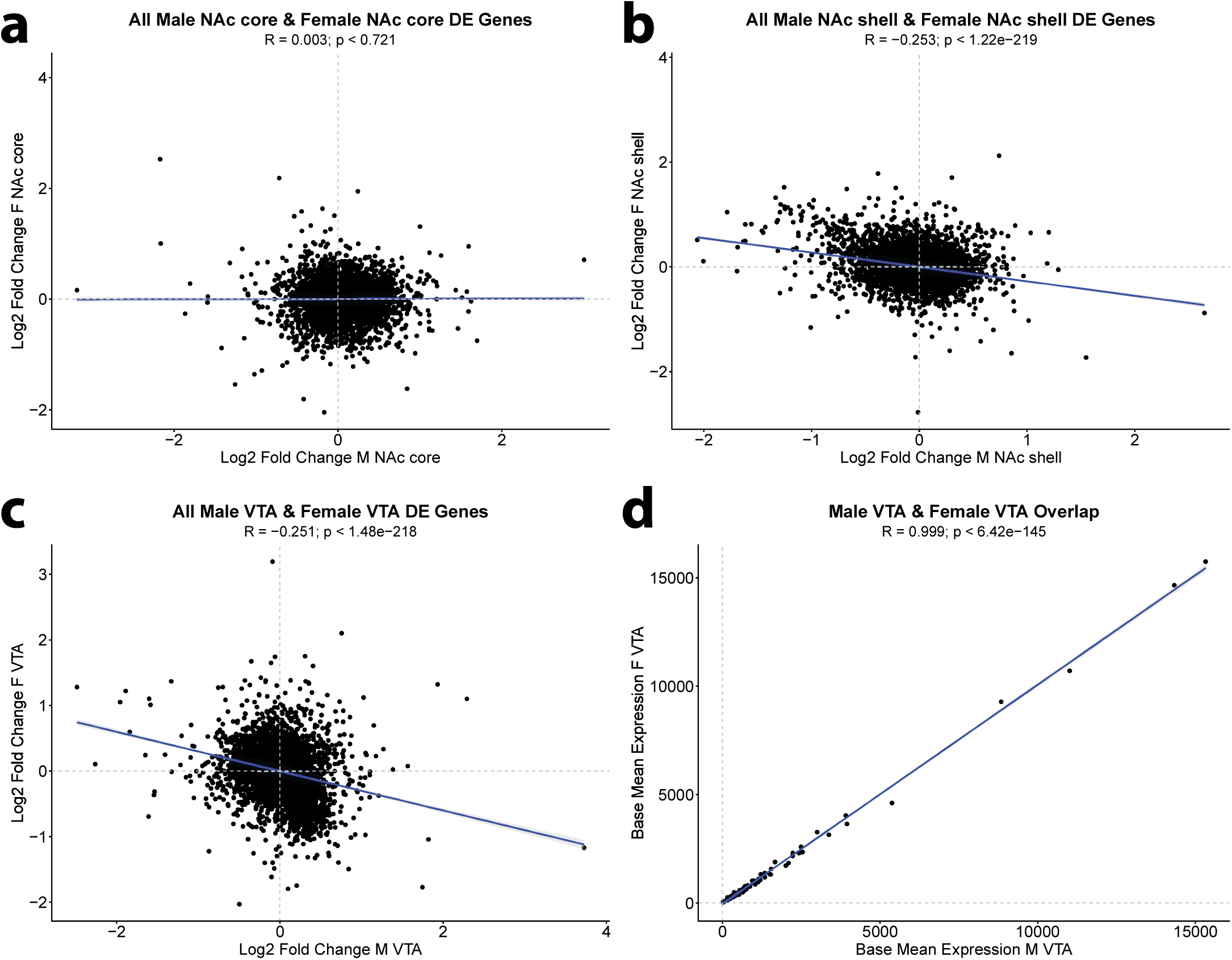
Transcriptome-wide correlation of gene expression fold-change between females and males in each brain region. (**a**) Log2-fold change in NAc core, (**b**) Log2-fold change in NAc shell, and (**c**) Log2-fold change in VTA. (**d**) Correlation of base mean expression (DESeq normalized count) in the male and female VTA for DE genes (with *p*<0.05 as cut-off).

**Extended Data Fig. 6.**
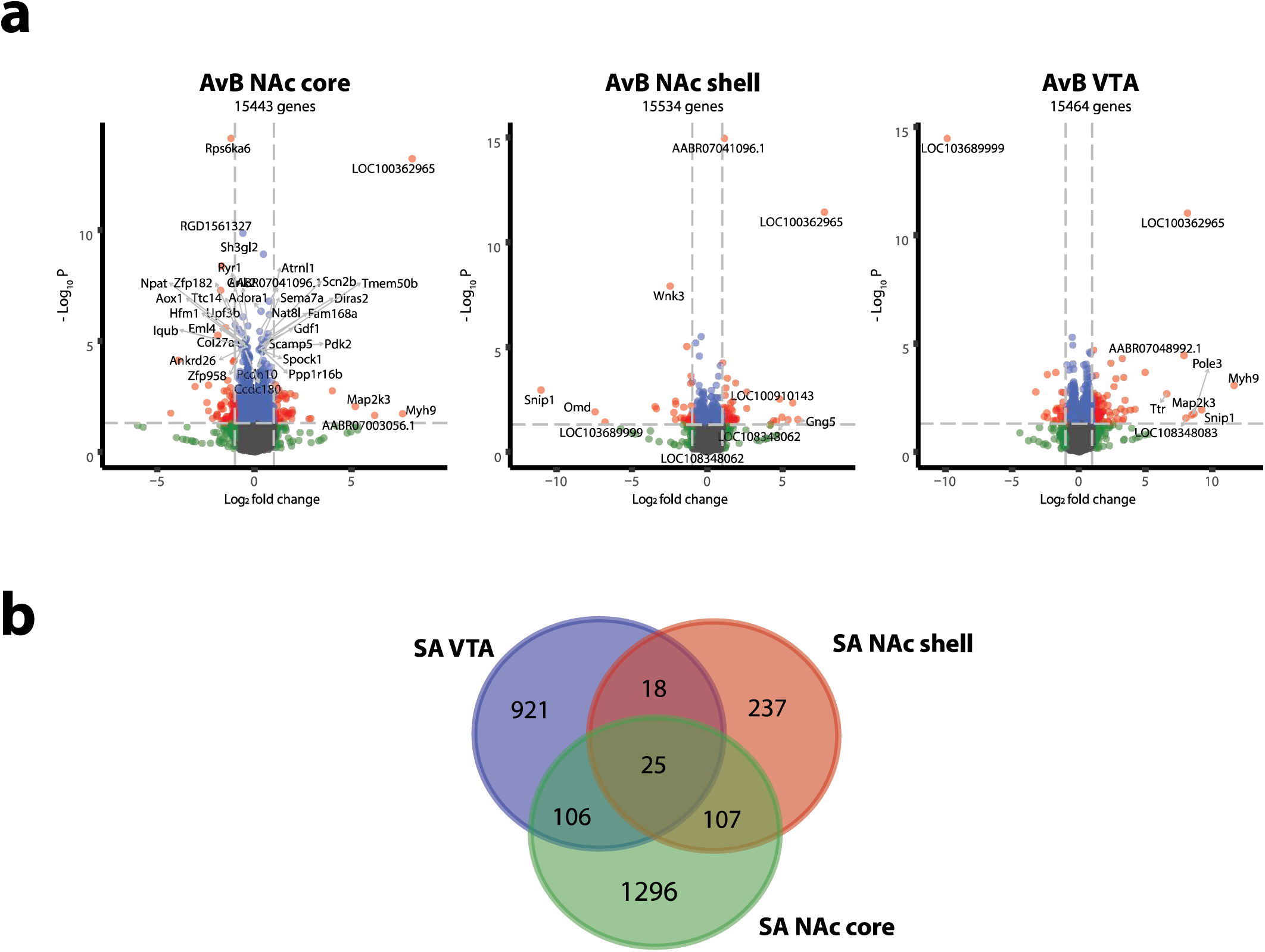
Differential expression (DE) of genes in tissues of self-administration A subgroup (SA-inclined) vs B subgroup (SA-disinclined) animals. (**A**) Volcano plots of DE genes in each brain tissue. The horizontal dashed lines indicate 2-fold changes and the vertical dashed lines indicate the nominal significance *p*-value of 0.05. (**b**) Venn diagram showing the overlapping DE (*p*<0.05) genes associated with NIC SA between different brain regions.

**Extended Data Fig. 7.**
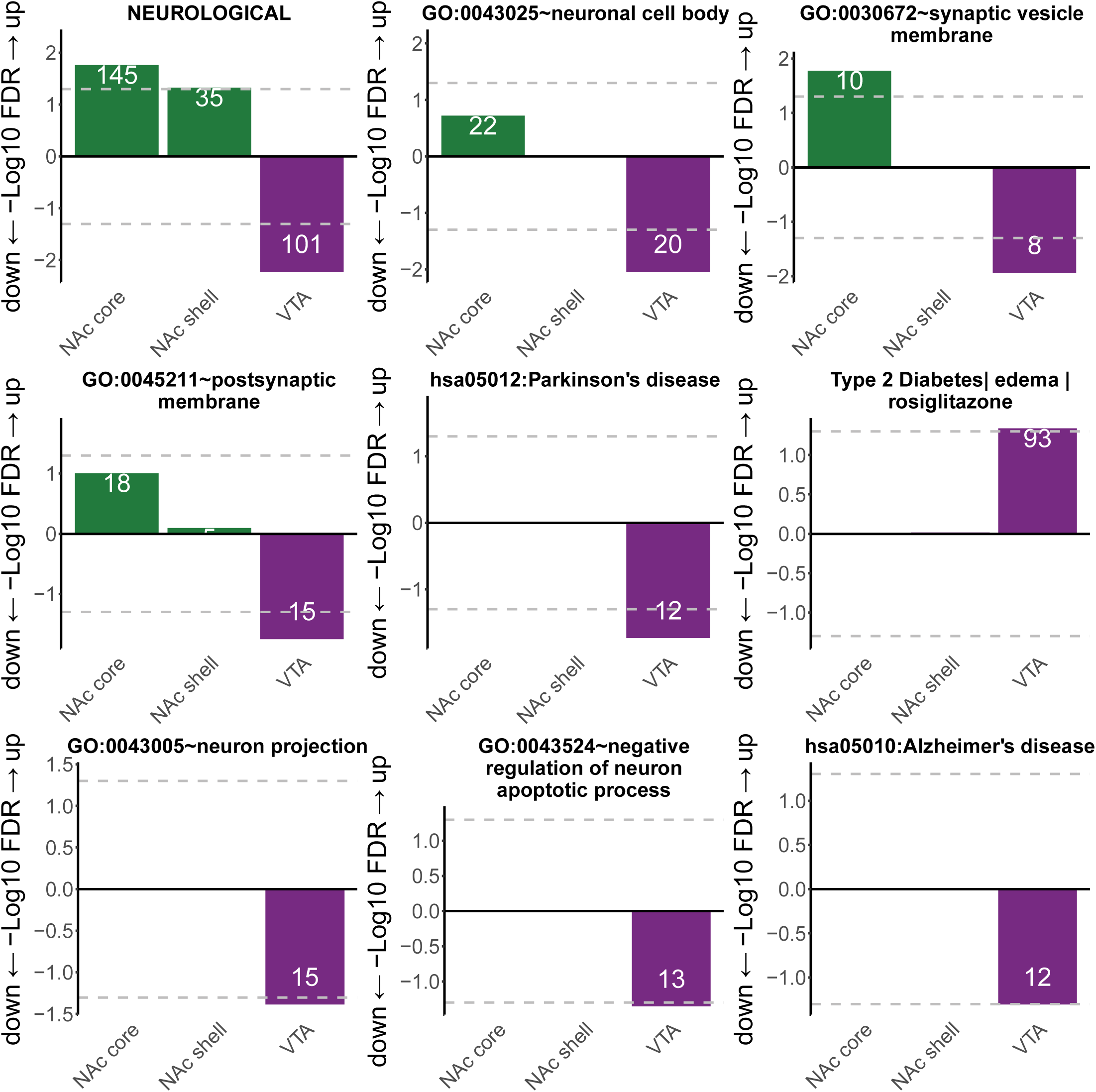
Extended DAVID gene set enrichment analysis of DE genes in each brain region for NIC self-administration. Listed are for GAD diseases and disease classes, OMIM diseases, KEGG pathways and GO terms. FDR-significant gene sets include the number of class genes (inset in the bar).

**Extended Data Fig. 8.**
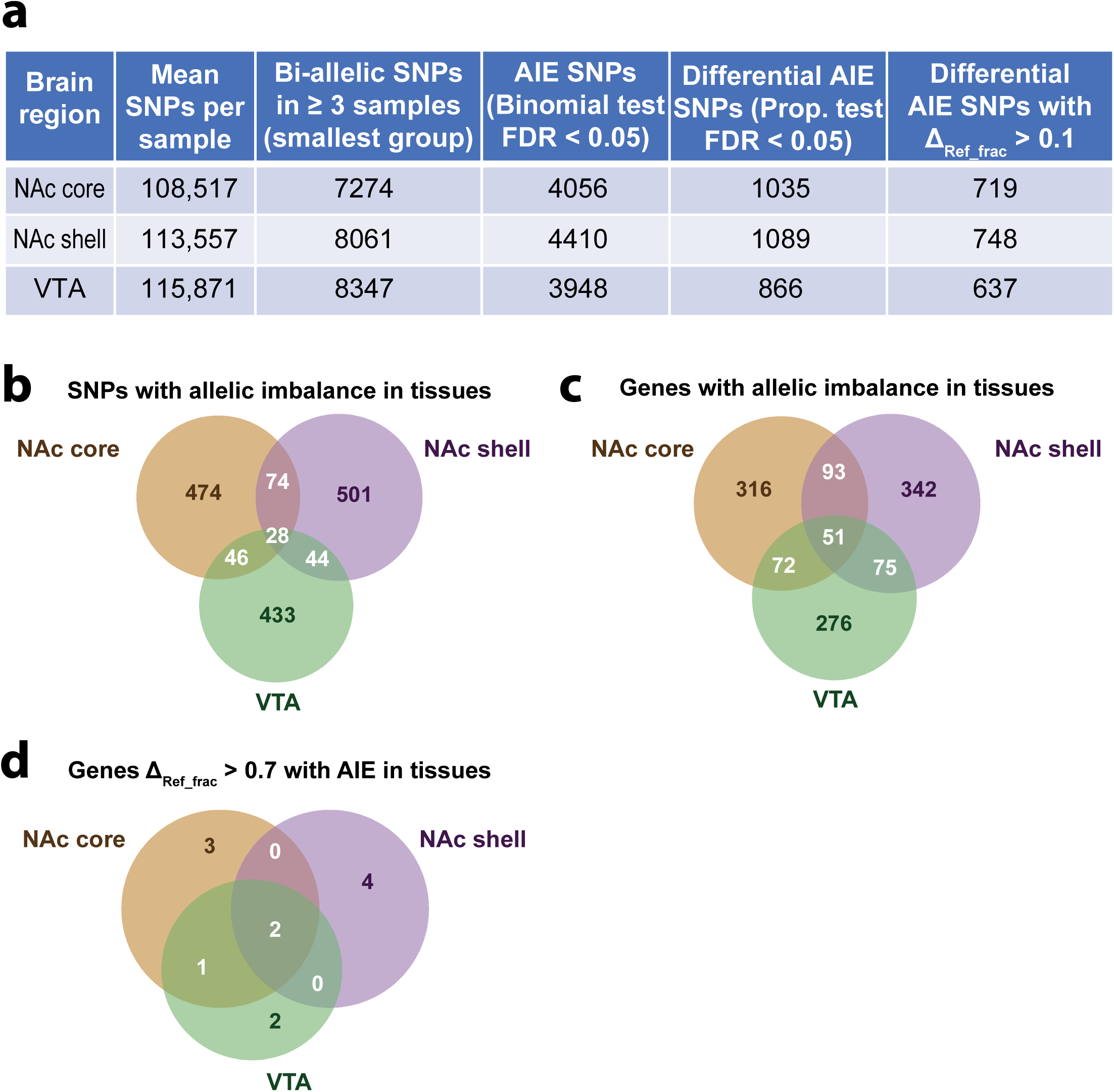
Allelic imbalance of expression (AIE) statistics in self-administration A subgroup (SA-inclined) and B subgroup (SA-disinclined) animals. (**a**) Total number of AIE SNPs at different cut-offs in each brain region. (**b**) Venn diagram of AIE SNPs in each brain region. (**c**) Venn diagram of genes containing AIE SNPs in each brain region. (**d**) Genes containing AIE SNPs with a relatively large difference of AIE between subgroups A and B (Reference allele fraction difference between subgroups>0.7).

**Extended Data Fig. 9.**
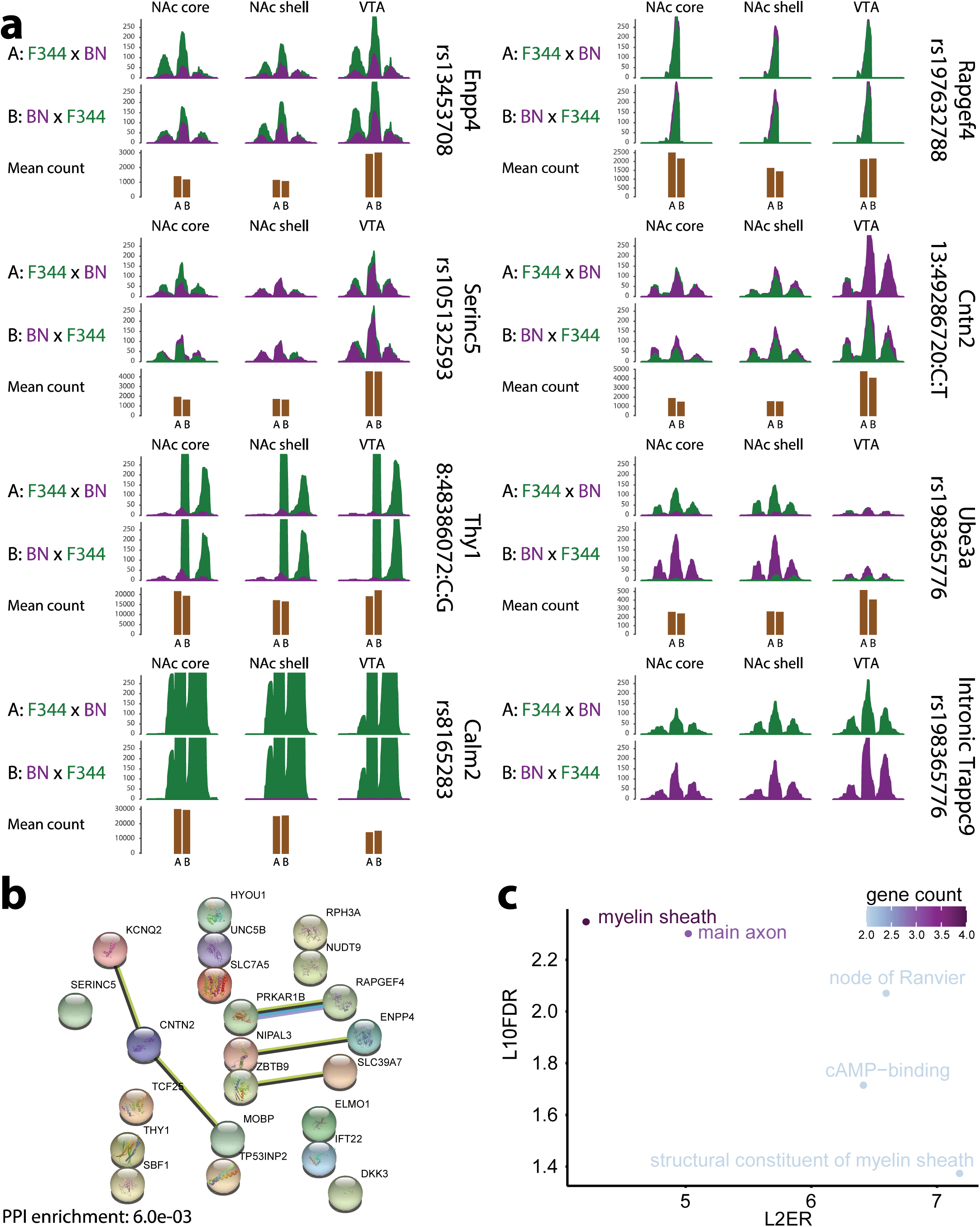
Allelic imbalance of expression (AIE) in genes with upregulated expression in NAc during self-administration (SA). (**a**) RNA-seq read pileup plots of example loci (transcribed SNPs) showing AIE in each brain region. The sequencing reads of the Fischer-344 (F344) allele are in green and those from the Brown Norway (BN) allele are in purple. Upper row depicts AIE of A subgroup (SA inclined), middle row depicts AIE of B subgroup (SA disinclined), and lower row depicts the normalized gene expression (mean RNA-seq read count) in A and B subgroups (shown in brown). (**b**) STRING analysis of human orthologs of the 22 genes with AIE and DE combined in the NAc core; with decreasing reference allele fraction in A subgroup And upregulated expression during SA (*p*<0.05). Number of nodes: 22, number of edges: 5, average node degree: 0.455, avg. local clustering coefficient: 0.364, expected number of edges: 1, PPI enrichment *p*<6.0×10^−3^. (**c**) Gene ontological enrichments from the STRING analysis in (**b**) colored by gene count in each term. X-axis indicates the Log2 enrichment ratio (L2RE) and Y-axis indicates the -Log10 FDR (L10FDR).

**Extended Data Fig. 10.**
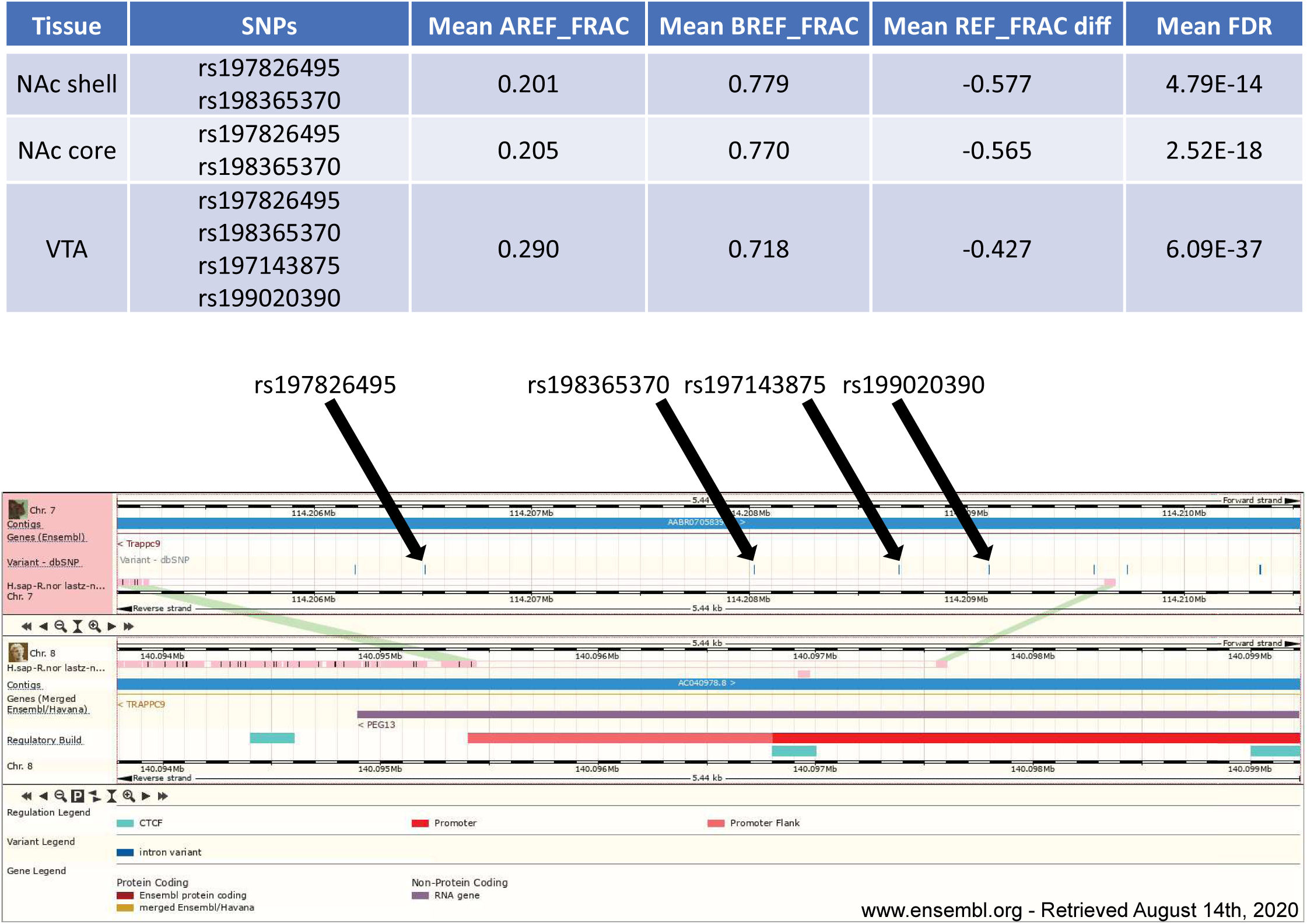
Intronic allelic imbalance of expression (AIE) SNPs in *Trappc9*, aligned to putative site of *paternally expressed gene 13* (*PEG13*) rat ortholog. Four transcribed SNPs showing strong AIE (upper table) in the intronic region of *Trappc9* identified in a rat (Rnor_6.0) to human (GRCh38.p13) LASTz alignment (lower; light pink), via ensemble.org. Rat *Trappc9* intron is shown in maroon in the rat genome, Human *PEG13* and *TRAPPC9* are shown in purple and yellow, respectively, on the human genome.

## Acknowledgements

This study was supported by National Institutes of Health (NIH) grant DA041600 (to J.D.).

## Author contributions

A.K. performed the main experiments and R.R.B. carried out the data analyses. A.K. and R.R.B. wrote the manuscript. S.Z. helped with AIE analysis and H.Z. helped with RNA-seq experiment. T.U. and S.S. performed the behavioral experiments. A.R.S. and Z.P.P helped with interpretation of results and edited manuscripts. P.V. supervised the behavioral experiments and wrote the manuscript. J.D. conceived the study, supervised the experimental work and data analyses, and wrote the manuscript.

## Animals and crosses

F1 progeny were generated by crossing two inbred Envigo rat strains (Fischer-344 [F344] and Brown Norway [BN]). Initial cross (F1i) was performed by crossing male F344 and female BN. Reciprocal cross (F1r) was performed by crossing female F344 and male BN. The F344 father/BN mother initial (subgroup A) and BN father/F344 mother (subgroup B) reciprocal cross F1 rats were evenly distributed in all experimental groups. The proven male and female breeding pairs (F344/NHsd and BN/RijHsd) were purchased from Envigo. The breeding was carried out at the University of Chicago Animal Resources Center by animal facility staff. All experimental behavioral procedures were performed during the dark phase of the light cycle according to an approved Institutional Animal Care and Use Committee protocol.

## Behavioral tests

For SST, F1 males (n=5-6/group) and females (n=7/group) were randomly assigned to three groups and tested in early adulthood (∼250 g). Rats in the first group were administered 0.1 mg/kg of NIC (all base and IP), those in the second 0.4 mg/kg, and those in the third administered saline (SAL;1.0 ml/kg). Injections were given once daily for 4 days. On days 1 and 4, rats were placed in an open field immediately after the injection and their locomotion measured for 2 hours. Rats were then left undisturbed in their home cages for 2 weeks after which they were tested for locomotor SST. On this test, all F1 rats were administered one injection of NIC (0.4 mg/kg), immediately placed in the open fields and their locomotion measured for two hours. For the SST transcriptomic analysis, F1 males (n=16) and females (n=16) were each divided into 2 groups. In the first (n=8), rats were administered 0.4 mg/kg NIC and in the second (n=8), SAL every day for 4 days. Brain tissues (∼20mg) from NAc core, NAc shell, and VTA regions were harvested two weeks later as described below. The open fields used to measure locomotion and the associated software were as described in Singer et al ^1^.

For NIC SA, F1 males (n=8) were prepared with an IV catheter and given the opportunity to self-administer NIC (30 μg/kg/infusion, base, IV) on each of 6 daily 2-hr self-administration hof 6daily 2 -hr self-administration sessions. Each self-administration chamber contained one lever, presses on which delivered a NIC infusion on a FR1 schedule of reinforcement. The number of actively self-administered infusions was recorded. Passive experimenter delivered IV priming infusions were administered at the start of each session and again following each 12-minute period of no active lever pressing. A maximum of 10 infusions (active + passive) was allowed per session. Thus, over the course of the six sessions, these rats received 60 infusions (active + passive) or 1.8 mg/kg NIC base, roughly equivalent to the 1.6 mg/kg administered in the locomotor sensitization experiments. To control for non-drug lever pressing, a non-catheterized group of no NIC self-administration control F1 rats was also tested (n=8). All remaining apparatus configurations and procedures were as described earlier ^2^. For the SA transcriptomic analysis, brain tissues from the NAc core, NAc shell, and VTA were harvested 3 days following the last session as described below.

## Brain tissue harvesting and homogenization

Brain tissues were hand dissected as described in Singer et al ^3^. Briefly, tissues were obtained bilaterally from 2mm coronal sections (VTA: -6.84 to -4.80; NAc: +1.08 to +3.00; mm from bregma ^4^) and the hemispheres combined for each site. 350 µl of cold lysis buffer (20 mM Tris, pH 7.4; 150 mM NaCl; 5mM MgCl_2_; 1mM DTT; 1% Triton X-100) was added to the brain tissue and sonicated (5 rounds: 3 sec – pulse, 12 sec – pause, amp 7). Tissue was triturated ten times through a 26-gauge needle (VWR, BD305111) and passed through Qiagen shredder (Qiagen, 79654) for 2 min at 14,000 × rpm at 4°C.

## RNA isolation and RNA sequencing (RNA-seq)

Total RNA was isolated from 30 μl of lysate using TRIzol LS reagent (Life technologies, 10296028). Lysate volume was adjusted to 100 μl and 300 μl Trizol LS (3× volume) was added. Then sample was incubated for 5 min at room temperature and 80 μl /(15volume)of chloroform was added following by shaking for 15 sec, incubation for 10 min at room temperature, and centrifugation for 15 min at 14,000 × rpm at 4°C. The aqueous phase containing the RNA was transferred to a new tube, 200 μl (1× volume) of isopropanol (VWR, 87000–048) was added to the aqueous phase and mixed well by inverting several times following by incubation for 10 min at room temperature and centrifugation for 25 min at 14,000 × rpm at 4°C. Total RNA precipitate formed a white gel-like pellet at the bottom of the tube. The pellet was washed with 1 ml of 75% ethanol and centrifuged for 5 min at 7,500 × rpm at 4°C. The pellet was washed one more time in 1 ml of 100% ethanol and centrifuged again at the same settings. The RNA pellet was dried for 10 min at room temperature and resuspended in 30 μl of RNase-free water. The RNA sample was cleaned with RNeasy MinElute Cleanup Kit (Qiagen, 74204). Total RNAs extracted from each sample were sent to the University of Minnesota Genomic Center for library preparation and RNA-seq with standard QC metrics (RNA quantity > 1 µg, RIN > 9.0, OD260/280 > 1.9), NovaSeq S2 2×50bp, > 32M PE reads per library.

## Gene expression analysis by qPCR

For qPCR, reverse-transcription was performed using ThermoFisher High-capacity RNA-to-cDNA reverse transcription kit (Applied Biosystems, 4366596) with random hexamers according to the manufacturer’s protocol. Briefly, 150-300 ng of total RNA was used for each 20 µl RT reaction. The reaction products were then diluted with 80 µl of RNase-free water for qPCR analysis. qPCR was performed using TaqMan Universal PCR Master Mix (Applied Biosystems, 4364338) on a Roche 480 II instrument, using gene-specific FAM-labelled TaqMan probes (ThermoFisher) for detecting gene expression, with GAPDH as the internal control. 7 biological replicates (male VTA brain tissues to test SST associated genes and NAc core brain tissues to test SA associated genes) were included for each experimental group, and 3 technical replicates were included in qPCR. Student’s *t-*test was used to quantify statistical significance in differential gene expression for NIC vs SAL (for SST associated genes *Klhdc8b, Ctnna2, Fbx117*) and subgroup A vs subgroup B rats (for SA associated genes *Rev3l, Fam168a, Cttnbp2*).

## Transcriptomic Analysis

RNA-seq reads were trimmed with fastp 0.20.0 ^5^ using -3, -5, and -l 20 settings and trimming for custom sequencing adapters. Transcriptional abundance was calculated from Rnor version 6 ^6^ with salmon 0.12.0 ^7^ with recommended settings for paired reads and collapsed to gene counts using tximport ^8^. SST samples had 71-77% read mapping, and SA samples had 72-76% read mapping. Principal component analysis (PCA) and DE were performed by using DEseq2 1.26.0 ^9^ with recommended parameters ^10^. Genes were limited to protein-coding and long non-coding RNA gene biotypes with biomaRt 2.42.0 ^11^. A total of 16,069 genes and 16,063 genes had detectable expression (normalized reads greater than 10 in at least 3 samples) for the SST and SA experiments, respectively. PCA utilized the variance stabilizing transformation of DESeq normalized counts ^10^. One outlier for male NAc core in SST and one for subgroup A group VTA in SA were discarded as extreme outliers based on PCA analysis. Pairwise sample distances were calculated from raw gene counts using Poisson distance with PoiClaClu 1.0.2.1 ^12^. Gene information and rat-human ortholog matching was conducted with biomaRt ^11^ using Ensembl version 99 ^13^ (which maps to human genome GRCh38.p13).

## Allelic Imbalance of Expression (AIE) Analysis

For the SA data set, allelic imbalance was calculated using Rnor version 6 ^6^ as the reference genome. To mitigate potential mapping bias to reference allele, correction for mapping bias was done by masked fasta. Using the merged reads of 2 samples of each F1 type and tissue (12 total samples, 2 each from subgroup A NAc core, subgroup A NAc shell, subgroup A VTA, subgroup B NAc core, subgroup B NAc shell, subgroup B VTA), samples were merged and mapped to reference using STAR ^14^ and SNPs were called via GATK 4.1.4.1 ^15^. SNP calls from merged reads were used to generate a masked fasta reference sequence with BEDTools 2.27.1 ^16^, which was then used to re-map using STAR. The resulting bam files were run through GATK HaplotypeCaller in gvcf format and combined with GenotypeGVCFs to generate a multi-sample all-calls vcf file. Bi-allelic SNPs were filtered for a minimum depth of 20 reads in all samples, ≥ 2 reference reads, and ≥ 2 alternate reads in at least 3 samples to exclude *de novo* and somatic SNP calls. Allelic depths in A and B groups were summed across samples, and flagged as imbalanced in subgroup A or subgroup B by binomial test with an FDR < 0.05, then flagged as phenotype relevant with the two-sample proportion test comparing the subgroup A and subgroup B groups with an FDR < 0.05 (two-tailed test). Final thresholding of biological relevance required a difference in reference allele (BN copy) fraction between subgroup A and subgroup B groups to be > 0.1. SNPs passing filtering were then annotated by gene location retaining only exonic and splice junction (±3bp) SNPs.

## Gene enrichment Analysis

Gene sets associated with NIC addiction were derived from results of Liu et al ^17^ for age of initiation (AOI), smoking initiation (SIn), smoking cessation (SCe), cigarettes per day (CPD), and drinks per week (DPW). Genes harboring risk variants within linkage disequilibrium (r^2^ > 0.3) for each NIC addiction phenotype ^17^ were checked for enrichment in SST and SA DE gene sets by Fisher’s exact test against a background of all expressed genes. SST and SA DE genes were also tested via Fisher’s exact test for enrichment of chromosome X genes. For enrichment of AIE genes, we used Fisher’s exact test against a background of all protein-coding and long non-coding RNA genes (the same constraint used above for transcriptomic analysis). SA AIE genes were tested for enrichment of NIC addiction phenotypes, chromosome X, and SA DE genes.

Human orthologs of SST and SA genes were tested for enrichment of GAD disease phenotypes ^18^, KEGG Pathways ^19^, and GO enrichments using DAVID ^20^. Finally, STRING protein interaction network analysis ^21^ was conducted on orthologs for interaction enrichments.

## GWAS enrichment test using MAGMA

MAGMA ^22^ enrichment of GWAS disease risk loci for SST and SA gene sets was done as previously described ^23^. Enrichment was tested on a range of upstream and downstream gene annotation windows (notable enrichments in the results section were present using multiple annotation windows), with 100kb upstream/downstream reported. Preliminary analysis of covariates was done to identify confounding variables from gene biotypes, and the HLA region. Covariates included in the final MAGMA model were expressed genes, protein-coding genes, and genes in the HLA-region (6:25000000-34000000). Summary statistics were from GWAS data sets of NIC phenotypes ^17^ and BMI ^24^. GWAS SNPs were limited to MAF > 0.01 in the Haplotype Reference Consortium r1.1 ^25^, and minimum INFO score > 0.9 in any trait with available data (at least one INFO score per SNP). SNPs were converted to GRCh38 using bcftools 1.9 ^26^ and custom scripts (see data availability).

## Statistical analyses

Behavioral data were analyzed by between and between-within analyses of variance (ANOVA) followed by post hoc Scheffé comparisons using the IBM SPSS Statistics module. The statistical analyses in DE test and gene set enrichment analyses were indicated separately in each section.

## Data Availability

All code used to generate results for this analysis is available at: https://rbutleriii.github.io/center_for_psychiatric_genomics/

Sequence data and raw count matrices have been submitted to the Gene Expression Omnibus for SST and SA experiments, GSE157726 and GSE157683, respectively.

---

**Supplementary Figure 1.**
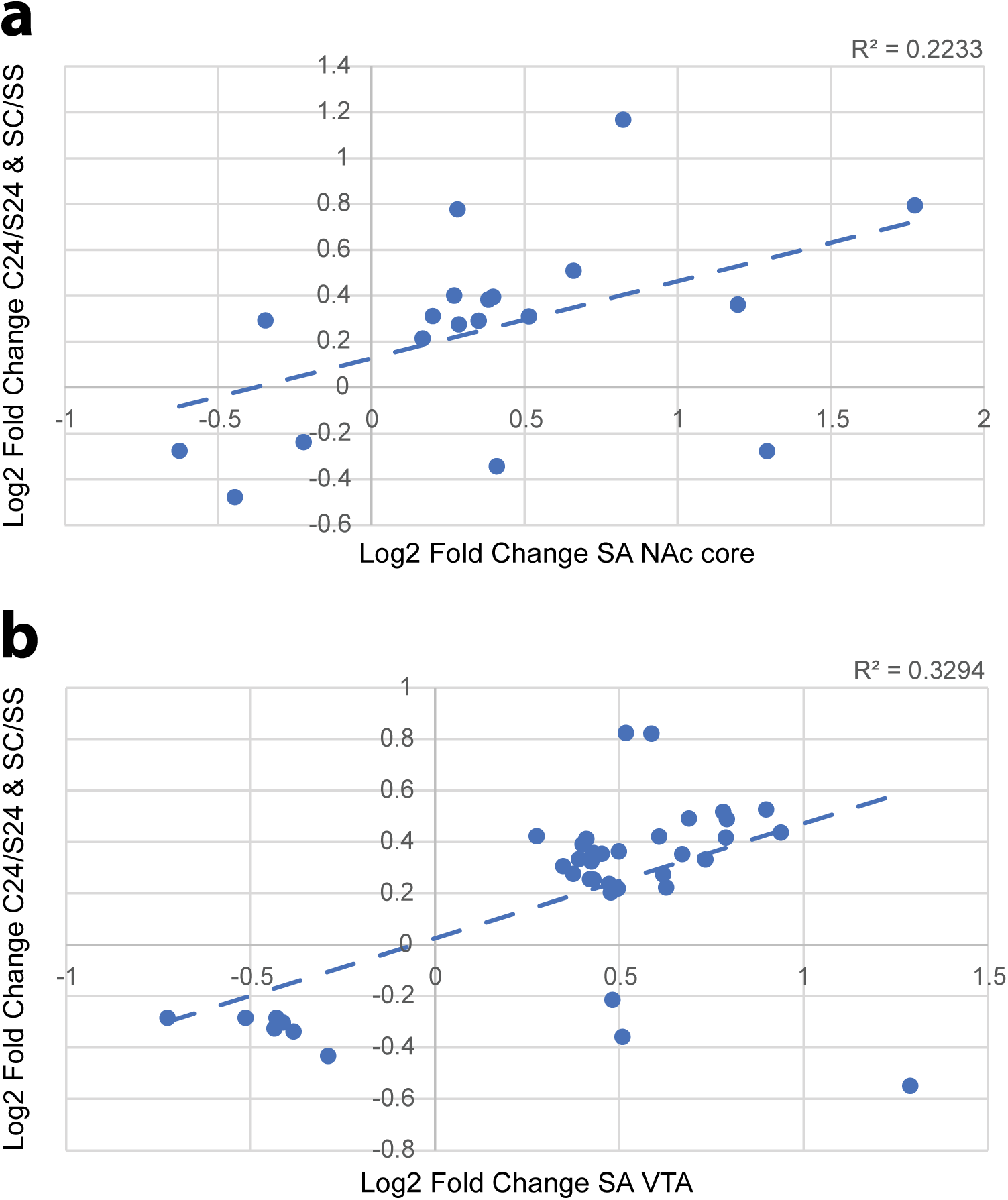
Correlation between self-administration (SA) gene differential expression (DE) and cocaine addiction gene DE in Walker et al 2018. **(A)** DE of SA nucleus accumbens (NAc) core genes that are also significantly DE in cocaine SA + 24-hour withdrawal (C24) vs saline SA + 24-hour withdrawal (S24) and saline SA + 30-day withdrawal + cocaine exposure (SC) vs saline SA + 30-day withdrawal + saline exposure (SS). **(B)** DE of SA ventral tegmental area (VTA) genes that are also significantly DE in C24 vs S24 and SC vs SS.

