## Supplementary Tables 1-14 for "Transcriptomic signatures of sex-specific nicotine sensitization and imprinting of self-administration in rats inform GWAS findings on human addiction phenotypes": 20200918_Supplementary_Figure_1.pdf

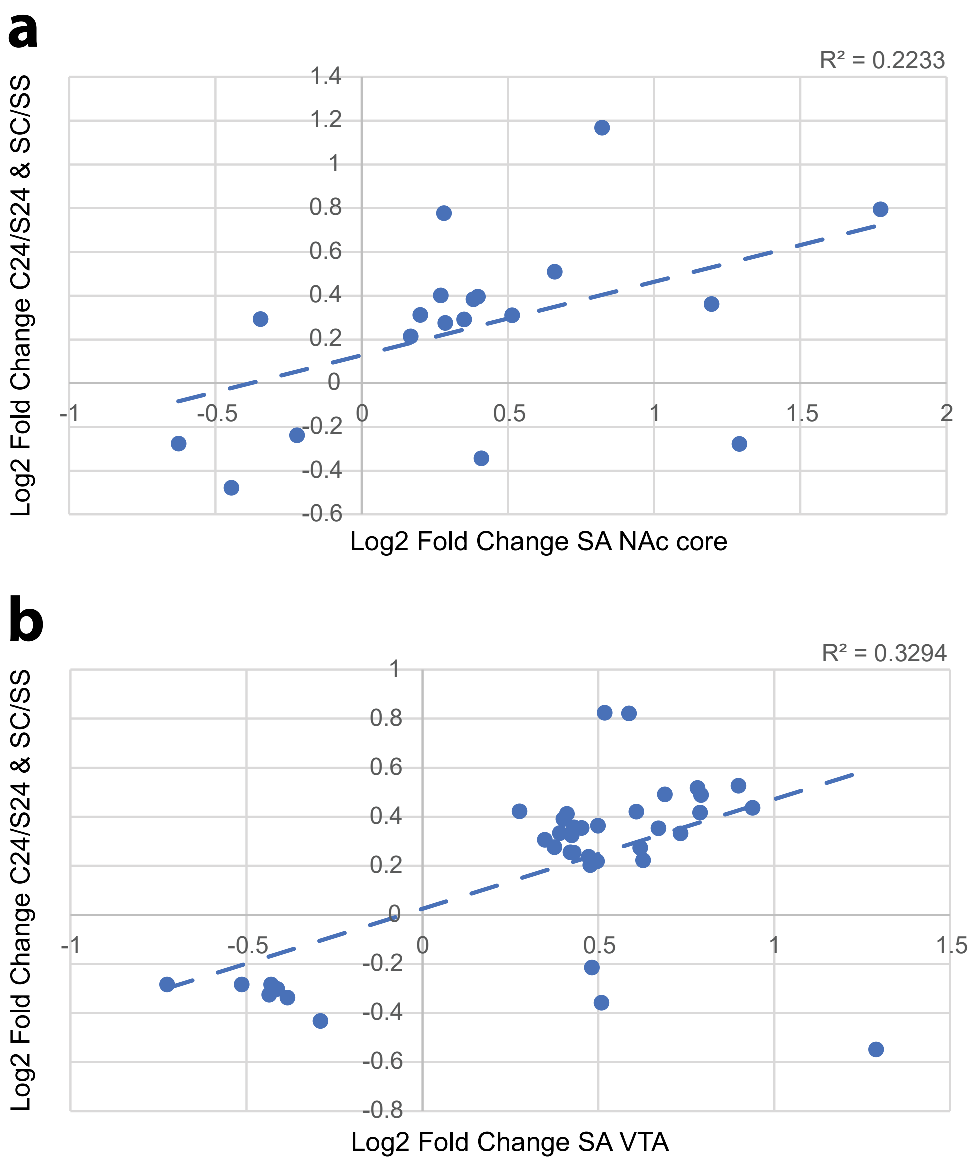


Supplementary Figure 1. **Correlation between self-administration (SA) gene differential expression (DE) and cocaine addiction gene DE in Walker et al 2018. (A)** DE of SA nucleus accumbens (NAc) core genes that are also significantly DE in cocaine SA + 24-hour withdrawal (C24) vs saline SA + 24-hour withdrawal (S24) and saline SA + 30-day withdrawal + cocaine exposure (SC) vs saline SA + 30-day withdrawal + saline exposure (SS). **(B)** DE of SA ventral tegmental area (VTA) genes that are also significantly DE in C24 vs S24 and SC vs SS.
